# Neuronal octopamine signaling regulates mating-induced germline stem cell proliferation in female *Drosophila melanogaster*

**DOI:** 10.1101/2020.03.20.999938

**Authors:** Yuto Yoshinari, Tomotsune Ameku, Shu Kondo, Hiromu Tanimoto, Takayuki Kuraishi, Yuko Shimada-Niwa, Ryusuke Niwa

## Abstract

Stem cells fuel the development and maintenance of tissues. Many studies have addressed how local signals from neighboring niche cells regulate stem cell identity and their proliferative potential. However, the regulation of stem cells by tissue-extrinsic signals in response to external cues remains poorly understood. Here we report that efferent octopaminergic neurons projecting to the ovary are essential for germline stem cell (GSC) proliferation in response to mating in female *Drosophila*. The neuronal activity of the octopaminergic neurons is required for mating-induced GSC proliferation as they relay the mating signal from Sex peptide receptor-positive cholinergic neurons. Octopamine and its receptor Oamb are also required for mating-induced GSC proliferation via intracellular Ca^2+^ signaling. Moreover, we identified Matrix metalloproteinase-2 as a downstream component of the octopamine-Ca^2+^ signaling to induce GSC proliferation. Our study provides a mechanism describing how neuronal system couples stem cell behavior to external cues through stem cell niche signaling.

## Introduction

Animal tissues are built from cells originally derived from stem cells (Spradling et al., 2001). During normal development and physiology, this robust stem cell system is precisely regulated (Drummond-Barbosa, 2008). Conversely, the dysregulation of these cells can result in abnormal tissue integrity and lead to deleterious diseases. Previous studies have revealed that many types of stem cells reside in a specialized microenvironment, or niche, where they are exposed to local signals required for their function and identity (Morrison and Spradling, 2008; Spradling et al., 2001). Recently, researchers have demonstrated how stem cell activity is regulated by tissue-extrinsic signals, such as hormones and neurotransmitters. For instance, in mammals, hematopoietic stem cells, mammary stem cells, muscle stem cells, and neural stem cells are influenced by sex hormones such as estrogen (Asselin-Labat et al., 2010; Bramble et al., 2019; Kim et al., 2016; Nakada et al., 2014). Retinoic acid and thyroid hormone play essential roles in the differentiation of testicular stem cells and neural stem cells, respectively (Gothié et al., 2017; Ikami et al., 2015). In addition, mesenchymal stem cell proliferation is stimulated by adrenaline (Wu et al., 2014). However, the when, how, and why these humoral factors are produced, circulated, and received during stem cell regulation remain to be elucidated.

The ovaries of the fruit fly *Drosophila melanogaster* are an excellent model system on how stem cell lineages are shaped by both local niche signals and tissue-extrinsic signals (Drummond-Barbosa, 2019). *D. melanogaster* ovary is composed of 16–20 chains of developing egg chambers called ovarioles. The anterior-most region of which, known as the germarium, contains germline stem cells (GSCs) that give rise to the eggs (Figure 1A and B). GSCs are adjacent to the somatic niche cells, which comprises cap cells, escort cells, and terminal filament cells (Figure 1A). After GSC divides, one daughter cell that remains attached to the niche cells retains its GSC identity, whereas the remaining daughter cell is displaced away from the niche cells and differentiates into cystoblast (CB). Each CB then undergoes differentiation into 15 nurse cells and 1 oocyte in each egg chamber, which is surrounded by somatic follicle cells.

**Figure 1.**
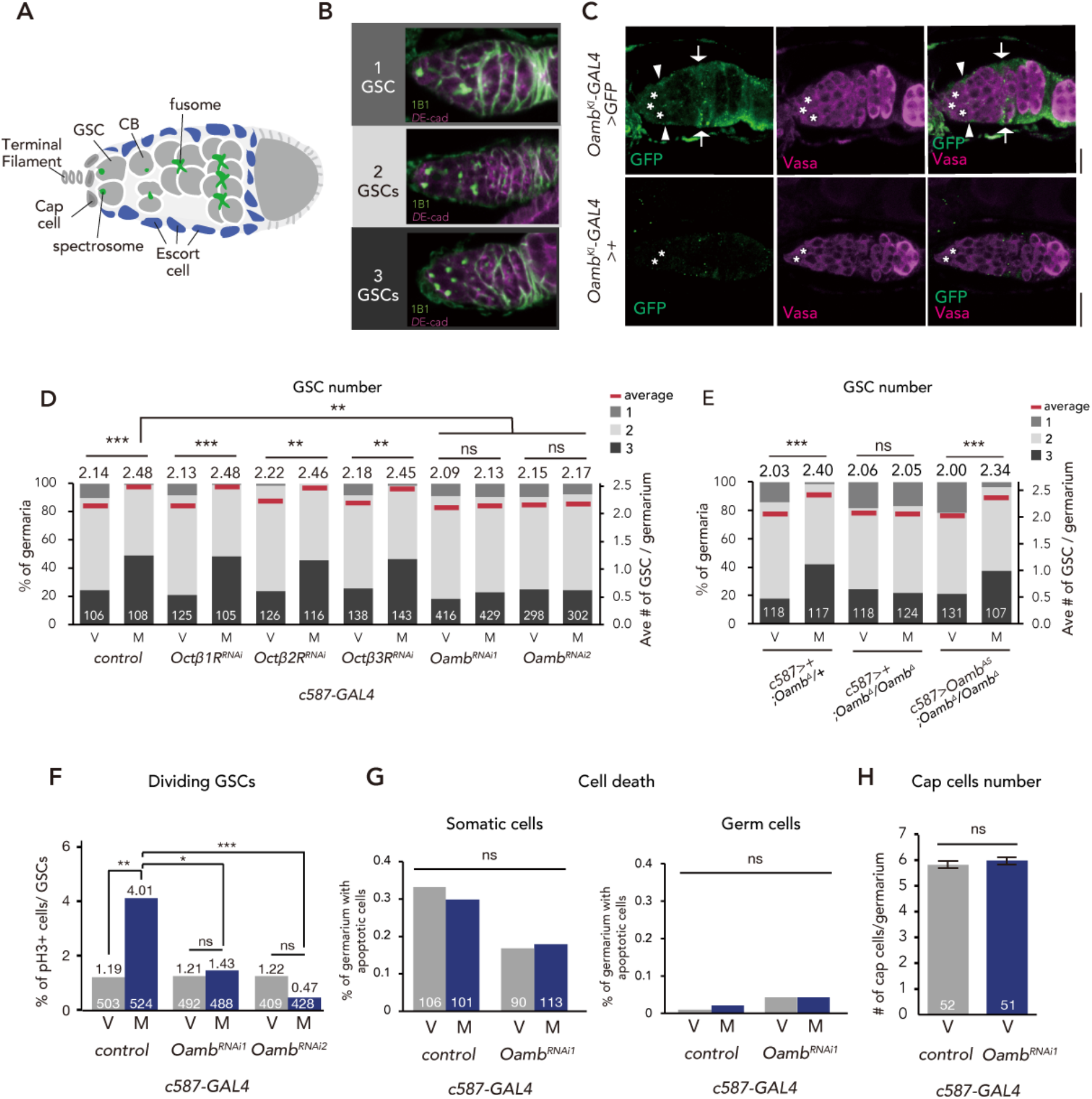
Post-mating GSC increase requires Oamb in the escort cells. (A) A schematic representation of *Drosophila* germarium. GSCs reside in a niche consisting of somatic cells such as cap cells, terminal filament cells, and escort cells and are identifiable by their stereotypical spectrosome morphology and location (adjacent to cap cells). GSC division produces 1 self-renewing daughter and 1 cystoblast (CB) that differentiates into a germline cyst. (B) Representative images of wild-type (*w^1118^*) female adults, containing 1, 2 and 3 GSCs from top to bottom. The samples were stained with monoclonal antibody 1B1 (green) and anti-DE-cadherin (magenta), which stain the spectrosome and overall cell membranes, respectively. (C) Immunofluorescence of germarium in adult female flies expressing the *20xUAS-6xGFP* reporter under *Oamb^KI^-GAL4*. *GFP*-expressing escort cells (arrowheads) are adjacent to GSCs (asterisk) and germ cells (magenta). GFP signal is also detected in follicle cells (arrow). Note that green fluorescence signals in escort cells are not observed in the germarium of *Oamb^KI^>+* female adults not expressing *GFP*. Scale bar 20 µm. (D-E) Frequencies of germaria containing 1, 2, and 3 GSCs (left vertical axis) and the average number of GSCs per germarium (right vertical axis) in virgin (V) and mated (M) female flies. *c587>+* flies were used as the control. (F) The ratio of pH3^+^ GSCs per total GSCs. (G) The ratio of apoptotic (Dcp-1^+^) somatic cells and germ cells per germarium. *c587>+* flies were used as the control. (H) The number of cap cells per germarium in the control and *Oamb* RNAi driven by *c587-GAL4*. Values on the y-axis are presented as the mean with standard error of the mean. *c587>+* flies were used as the control. For D-H, the number of germaria analyzed is indicated inside the bars. Wilcoxon rank sum test with Holm’s correction was used for D, E, and H. Fisher’s exact test with Holm’s correction was used for F and G. ***P ≤ 0.001, **P ≤ 0.01, and *P ≤ 0.05; NS, nonsignificant (P > 0.05).

GSC niche produces and secretes several local niche signals that regulate the balance between GSC self-renewal and differentiation (Hayashi et al., 2020; Kirilly and Xie, 2007; Spradling et al., 2011). For example, bone morphogenetic protein (BMP) ligands Decapentaplegic (Dpp) and Glass bottom boat (Gbb) are produced from the niche cells and directly activate BMP receptors in GSCs, leading to the repression of the differentiation inducer, *bag-of-marbles* (*bam*) (Morrison and Spradling, 2008; Zhang and Cai, 2020). Recent *D. melanogaster* GSC studies have also contributed to understanding of the systemic regulation of stem cell proliferation and maintenance in response to external environmental cues (Ables and Drummond-Barbosa, 2017; Drummond-Barbosa, 2019; Lin and Hsu, 2020; Yoshinari et al., 2019). For example, protein restriction results in a reduction in GSC division, which is mediated by *Drosophila* insulin-like peptides (DILPs) (LaFever and Drummond-Barbosa, 2005). In addition, nutrients influence GSC maintenance via the adipocyte metabolic pathway (Armstrong and Drummond-Barbosa, 2018; Matsuoka et al., 2017).

Besides nutrients, we have recently found out that mating is another external cue that significantly affects *D. melanogaster* GSC increase. Mated females show a dramatic increase in egg production, as well as GSC, which is induced by a male-derived peptide from the seminal fluid called Sex peptide (SP) (Kubli, 2003; Yoshinari et al., 2019). SP is received by its specific receptor, Sex peptide receptor (SPR), in a small subset of SPR-positive sensory neurons (SPSNs), which innervate the uterine lumen and send afferent processes into the tip of the abdominal ganglion (Häsemeyer et al., 2009; Yapici et al., 2008). The SP-SPR signaling in the SPSNs stimulates the biosynthesis of the ovarian insect steroid hormones (ecdysteroids), which play an essential role in mating-induced GSC increase (Ameku et al., 2017; Ameku and Niwa, 2016; Uryu et al., 2015). Because SPSNs do not directly innervate into the ovary, it is hypothesized that a signal and its signaling pathway are involved in bridging the gap between SPSNs and GSCs. However, it is still unclear how mating information is transmitted from SPSNs to GSCs at the molecular and cellular levels.

Here, we present a series of new findings that reveal a novel and fundamental neuronal mechanism connecting SPSNs and GSCs to regulate mating-induced GSC increase. We demonstrate that a small subset of neurons directly innervating into the ovary plays an indispensable role in regulating mating-induced GSC increase. These neurons produce the monoamine neurotransmitter, octopamine (OA), the insect equivalent of noradrenaline (Roeder, 2005). We also show that the neuronal activity of the OA-producing neurons is required for mating-induced GSC increase. Moreover, we find that the OA directory activates GSC increase through its receptor, octopamine receptor in mushroom body (Oamb), followed by Ca^2+^ signaling in the ovarian escort cells. Furthermore, OA/Oamb signaling requires the Matrix metalloprotease 2 (Mmp2) to activate GSC increase in the ovarian escort cells. Finally, we show that SPSNs relay the mating signal to the ovary-projecting OA neurons via nicotinic acetylcholine receptor signaling. Taken together, we propose a novel efferent neuronal pathway that transmits mating stimulus to the GSC to control stem cell number. Our study provides a mechanism describing how neuronal system couples stem cell behavior to external cues, such as mating, through stem cell niche signaling.

## Results

### Mating-induced GSC increase requires the octopamine receptor Oamb in ovarian escort cells

Previous studies have reported that *Oamb*, one of the G protein-coupled receptors of OA, plays an essential role in the ovulation of *D. melanogaster* (Deady and Sun, 2015; Lee et al., 2009, 2003). Although *Oamb* is widely expressed in the female reproductive system (Deady et al., 2017; Deady and Sun, 2015; Lee et al., 2003), including the oviduct and mature follicle cells of stage 14 egg chambers, its expression in the germarium and ovarian somatic cells of the early ovariole remains unexamined. To explore this, we generated the *Oamb* knock-in *T2A-GAL4* line (*Oamb^KI^-GAL4*) by In-Frame Fusion (Diao and White, 2012; Kondo et al., 2020), in which the *GAL4* activity is expected to more precisely recapture the endogenous *Oamb* expression. When we observed the *Oamb^KI^-GAL4*-driven GFP signal, we detected expressions not only in the oviduct and mature follicle cells as previously reported (Deady and Sun, 2015) (Figure 1-figure supplement 1A), but also in the germarium. Particularly, *Oamb^KI^-GAL4* was detected in the germarium, including escort cells and follicle cells (Figure 1C).

Because some of the *Oamb^KI^-GAL4*–positive cells were escort cells located near GSCs (Figure 1C), we investigated whether GSC proliferation or maintenance is regulated by *Oamb* expressed in the germarium. Therefore, we conducted transgenic RNAi experiment to knock-down *Oamb* gene using *c587-GAL4* driver, which is active in the escort cells of the germarium (Manseau et al., 1997). In control females, the mated ones exhibited an increase in GSC number as we have reported previously (Ameku and Niwa, 2016) (Figure 1D). In contrast, *c587-GAL4*–mediated *Oamb* knock-down (*c587>Oamb^RNAi^)* showed significantly impaired mating-induced GSC increase (Figure 1D). The phenotype was observed with two independent *UAS-Oamb-*RNAi strains (*Oamb^RNAi^* and *Oamb^RNAi2^*) (Figure 1D), each of which targeted a different region in the *Oamb* mRNA. The specificity of *Oamb* was also confirmed by the fact that the *c587-GAL4*–driven transgenic RNAi of other octopamine receptor genes (*Octβ1R*, *Octβ2R* and *Octβ3R*) (Ohhara et al., 2012) had no significant effect on the GSC number between virgin and mated females (Figure 1D).

In addition, *Oamb* RNAi in the escort cells from *GAL4* line *Traffic jam (tj)-GAL4* (Olivieri et al., 2010) (*tj>Oamb^RNAi^*) resulted in the failure of mating-induced GSC increase, whereas *Oamb* RNAi in the cap cells (*bab>Oamb^RNAi^*) and germ cells (*nos>Oamb^RNAi^*) had no effect on GSC increase (Figure 1-figure supplement 1B and C). These results were consistent with the spatial expression pattern of *Oamb* in the germarium as described above (Figure 1C and Figure 1-figure supplement 1A).

Notably, *c587-GAL4* and *tj-GAL4* are expressed not only in the escort cells but also in the nervous system (Ameku et al., 2018). Moreover, *Oamb* is expressed in the nervous system (Han et al., 1998). However, *Oamb* RNAi in the nervous system (*nSyb>Oamb^RNAi^*) did not affect the mating-induced GSC increase (Figure 1-figure supplement 1C), suggesting that the impairment of GSC increase of *c587>Oamb^RNAi^* or *tj>Oamb^RNAi^* is not due to the gene knock-down in neuronal cells but rather to that in escort cells.

To confirm the role of *Oamb* in mating-induced GSC increase, we generated *Oamb* complete loss-of-function genetic allele by Clustered Regularly Interspaced Short Palindromic Repeats (CRISPR)/CRISPR-associated protein 9 (Cas9) technology (Kondo and Ueda, 2013) (Figure 1-figure supplement 1D). Similar to *Oamb* RNAi females, *Oamb* homozygous mutant females (*c587>+; Oamb^Δ^/Oamb^Δ^*) did not exhibit mating-induced GSC increase (Figure 1E). In addition, the GSC increase of *Oamb^Δ^/Oamb^Δ^* was restored by overexpression of *Oamb* in the escort cells (*c587>Oamb^AS^; Oamb^Δ^/Oamb^Δ^*). These findings are all consistent with the idea that Oamb in escort cells modulates GSC increase after mating.

### Oamb in escort cells is required for GSC proliferation after mating

Because mating-induced GSC increase is accompanied by GSC division (Ameku and Niwa, 2016), We next examined whether Oamb in the escort cells is involved in GSC division after mating. We determined the number of GSCs during the M phase by staining using anti-phospho-Histone H3 (pH3) in control and *c587>Oamb^RNAi^* females. In control female flies, mating increased the frequency of GSCs (Figure 1F), whereas in *c587>Oamb^RNAi^* flies, this was not observed. We also monitored the fraction of apoptotic cells in the germarium by staining with anti-cleaved death caspase-1 (Dcp-1), a marker for apoptotic cells (Song et al., 1997). The number of apoptotic cells in the germarium did not change in *c587>Oamb^RNAi^* female flies compared with controls (Figure 1G), suggesting that Oamb activates GSC increase by pushing the cell cycle of GSCs and that the lack of mating-induced GSC increase in *Oamb* RNAi is not due to the enhancement of cell death.

Further, we determined the number of cap cells, which are critical components of the GSC niche (Xie and Spradling, 1998). *c587>Oamb^RNAi^* did not change the number of cap cells in virgin nor mated female flies, suggesting that *Oamb* knock-down does not affect the overall architecture of the niche (Figure 1H). Overall, Oamb in the escort cells plays a pivotal role in mating-induced GSC proliferation.

Oamb in mature follicle cells and in the oviduct has significant role in ovulation (Deady and Sun, 2015; Lee et al., 2009, 2003), and it is possible that these *Oamb* may indirectly induce GSC increase after mating. However, as *Oamb* RNAi in the stage-14 follicle cells by *R44E10-GAL4* (Deady and Sun, 2015) (*R44E10>Oamb^RNA^*^i^) nor in the common oviduct by *RS-GAL4* (Lee et al., 2003) (*RS>Oamb^RNA^*^i^) neither impaired mating-induced GSC increase (Figure 1-figure supplement 1E and F), we conclude it to be unlikely. These data support the idea that mating-induced GSC increase is independent from the ovulation process.

### OA administration is sufficient to induce GSC increase and BMP signaling in GSCs in an Oamb-dependent manner

To examine whether OA is received in the ovary but not in other organs to induce GSC increase, we cultured dissected virgin ovaries *ex vivo* with or without purified OA in the culture medium. After incubation for 12 h, the ovaries cultured with OA had more GSCs as compared to those without OA (Figure 2-figure supplement 1A). Whereas the minimal OA concentration to induce *ex vivo* GSC increase was 1 µM, hereafter we used 100 µM OA because the GSC number reached plateau with this concentration (Figure 2-figure supplement 1A). This OA-mediated *ex vivo* GSC increase was not observed in *c587>Oamb^RNAi^* virgin ovaries (Figure 2A).

**Figure 2.**
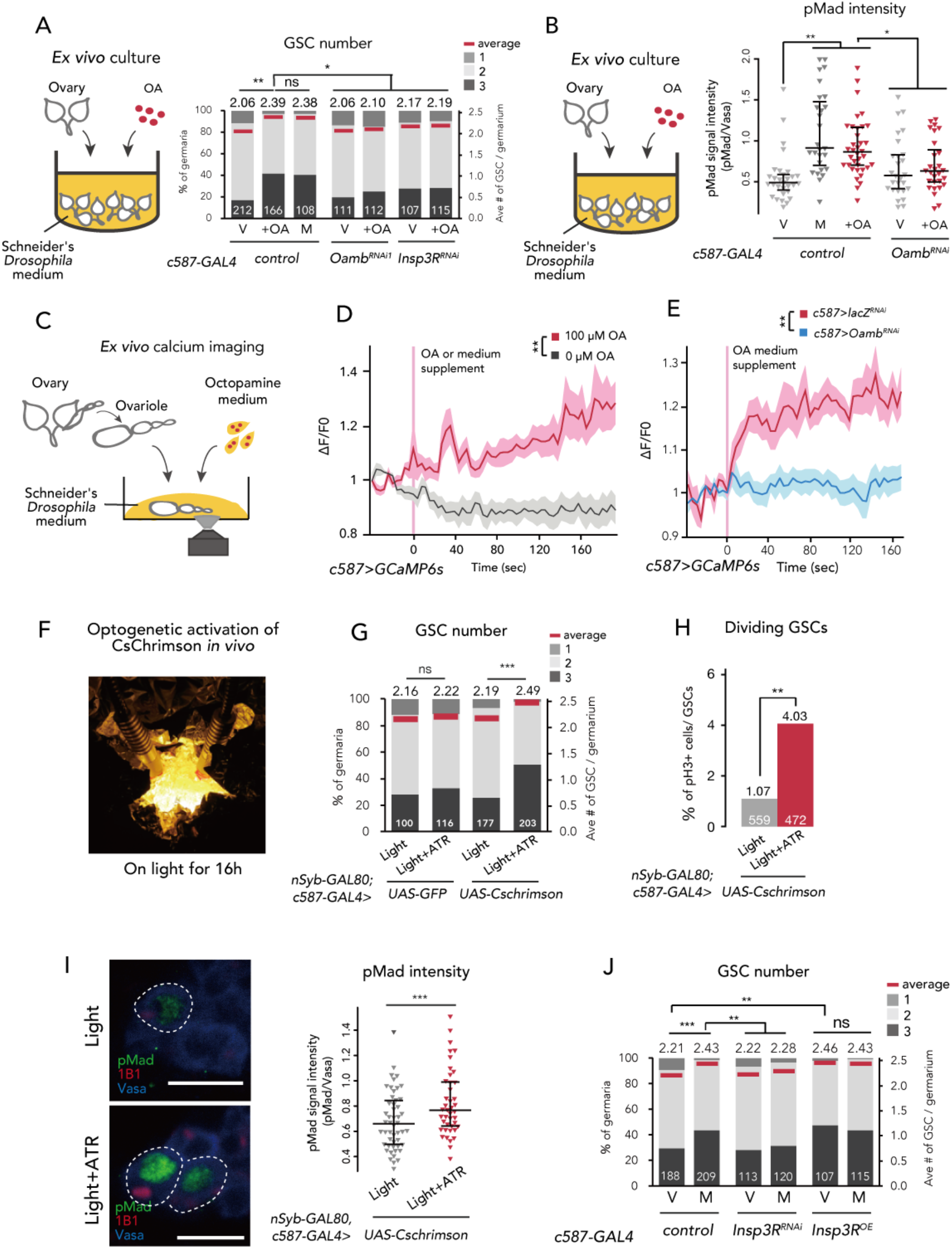
Ca^2+^ signaling is necessary for mating-induced GSC increase. (A) Frequencies of germaria containing 1, 2, and 3 GSCs (left vertical axis) and the average number of GSCs per germarium (right vertical axis). The ovaries were dissected from virgin (V), mated (M), and virgin ovaries cultured with OA (+OA). *c587>+* flies were used as the control. The number of germaria analyzed is indicated inside the bars. (B) Quantification of relative pMad intensity levels in the GSCs of *ex vivo* cultured ovaries (i.e., virgin (V), mated (M), and virgin cultured with OA (+OA)) as normalized to the pMad intensity in CBs. For the quantification of pMad intensity, the cell boundaries of GSCs and CBs were determined using anti-Vasa staining. Each sample number was at least 25. The 3 horizontal lines for each sample indicate lower, median, and upper quartiles. *c587>+* flies were used as the control. (C) A schematic representation of *ex vivo* calcium imaging. The dissected ovariole was incubated in Schneider’s *Drosophila* medium with or without OA. (D) Changes in the relative fluorescence intensity of GCaMP6s after 200 s without stimulation (n=8) or with stimulation (n=10) with 100 µM OA in *c587> GCaMP6s* Flies, and (E) with 100 µM OA as control (*c587>LacZ^RNAi^*, *GCaMP6s*; n=8) and *c587>Oamb^RNAi^*, *GCaMP6s* (n=8) female ovaries. Note that OA significantly increased the calcium response in escort cells, but *Oamb^RNAi^* impaired the calcium response. Statistical analysis was done at 120 s. (F) Equipment setup for optogenetic activation of CsChrimson. Flies were placed under the light for 16 h before dissection. (G) Frequencies of germaria containing 1, 2, and 3 GSCs (left vertical axis) and the average number of GSCs per germarium (right vertical axis) with light and with light and ATR in vrgin female. *nSyb-GAL80; c587>GFP* flies were used as control. The number of germaria analyzed is indicated inside the bars. (H) The ratio of pH3^+^ GSCs and total GSCs. The number of GSCs analyzed is indicated inside the bars. (I, left) Representative images of adult female germaria immunostained with anti-pMad antibody (green), anti-1B1 antibody (red), and anti-Vasa antibody (germ cell marker; blue) are shown. GSCs are outlined with dotted lines. (I, right) Quantification of the relative pMad intensity in GSCs, which was normalized to that in CBs. For the quantification of pMad intensity, the cell boundaries of GSCs and CBs were determined using anti-Vasa staining. Each sample number is at least 30. The 3 horizontal lines for each data sample indicate lower, median, and upper quartiles. (J) Frequencies of germaria containing 1, 2, and 3 GSCs (left vertical axis) and the average number of GSCs per germarium (right vertical axis) in virgin (V) and mated (M) female flies. *c587>+* flies were used as the control. The number of germaria analyzed is indicated inside the bars. Wilcoxon rank sum test with Holm’s correction was used for A, B, D, E, G, I, and J. Fisher’s exact test was used for H. ***P ≤ 0.001, **P ≤ 0.01, and *P ≤ 0.05; NS, non-significant (P > 0.05).

Our previous studies revealed that mating-induced GSC increase is mediated by GSC niche signals (Ameku et al., 2018; Ameku and Niwa, 2016). In particular, Dpp, the fly counterpart to BMP, is the essential niche signal (Spradling et al., 2011; Xie and Spradling, 1998). We therefore examined whether OA treatment affects Dpp signaling in GSCs *ex vivo* by measuring the level of phosphorylated Mad (pMad), a readout of Dpp signaling activation (Chen and McKearin, 2003; Raftery and Sutherland, 1999). We confirmed that OA treatment was sufficient to induce the increase in pMad level in GSCs, even in the *ex vivo* culture system (Figure 2B). On the other hand, OA treatment did not increase pMad levels in *c587>Oamb^RNAi^* ovaries (Figure 2B). These results suggest that OA activates the BMP signal in GSCs through Oamb in the escort cells, thereby resulting in the increase in GSCs.

### Ca^2+^ signaling in escort cells is essential for OA-dependent GSC increase

Upon OA binding, Oamb evokes Ca^2+^ release from the endoplasmic reticulum (ER) into the cytosol, leading to a transient increase in intercellular Ca^2+^ concentration ([Ca^2+^]_i_) (Han et al., 1998). To determine whether OA induces GSC increase via affecting [Ca^2+^]_i_ in escort cells, we first monitored [Ca^2+^]_i_ using a genetically encoded calcium sensor, GCaMP6s (Nakai et al., 2001; Ohkura et al., 2012). We dissected the virgin ovaries, in which *GCaMP6s* transgene was expressed driven by *tj-GAL4* or *c587-GAL4*, cultured them *ex vivo*, and then observed GCaMP6s fluorescence (Figure 2C). We found that 100 µM of OA treatment evoked an increase in GCaMP6s fluorescence in the escort cells and follicle cells, whereas the control medium treatment (0 µM of OA) did not show any increase in fluorescence (Figure 2D and Figure 2-figure supplement 1B; also see Video 1 and 2). In addition, the OA-mediated increase in GCaMP6s fluorescence intensity was not observed in the germarium of *Oamb* RNAi flies (Figure 2E), suggesting that the OA-dependent increase in [Ca^2+^]_i_ in the germarium is required for Oamb.

We also employed another approach utilizing the light-gated cation channel, CsChrimsonn (Klapoetke et al., 2014). We prepared *c587-GAL4*>*CsChrimson* flies in combination with *nSyb-GAL80* (Harris et al., 2015), allowing the expression of *CsChrimson* gene only in the germarim but not in the nervous system. Because CsChrimson requires all trans-retinal (ATR) to form its proper protein conformation (Wang et al., 2012), we utilized the flies fed with and without ATR supplement as experimental and control groups, respectively. When we irradiated an orange-light to *c587-GAL4, nSyb-GAL80*>*CsChrimson* flies to induce Ca^2+^ flux in the germarium (Figure 2F), GSC increased in the virgin females (Figure 2G). In addition, the ratio of GSC in the M phase increased by CsChrimson activation (Figure 2H). Moreover, the pMad level in GSCs was increased by CsChrimson activation, suggesting that the forced Ca^2+^ flux is sufficient to induce GSC increase through the upregulation of BMP signaling (Figure 2I).

We next confirmed whether the downstream component of Ca^2+^ signaling is involved in the mating-induced GSC increase. The knock-down of *Inositol 3-receptor* (*c587>Insp3R^RNAi^*) encoding a protein that releases the stored Ca^2+^ from ER suppressed GSC increase in mated females (Figure 2J). Conversely, the overexpression of *Insp3R* in the escort cells increased the GSC number in virgin females (Figure 2J). Furthermore, OA-mediated *ex vivo* GSC increase was not observed in *c587>Insp3R^RNAi^* virgin ovaries (Figure 2A). Overall, we demonstrated that OA signaling regulates the mating-induced GSC increase by controlling Ca^2+^ signaling in escort cells, which is thereby necessary and sufficient to induce GSC proliferation.

### Ovarian ecdysteroid signaling is required in the OA-Oamb-Ca^2+^-dependent GSC increase

The biosynthesis and signaling of ecdysteroid in the ovary are required for the mating-induced GSC increase (Ameku et al., 2017; Ameku and Niwa, 2016). Therefore, we next examined whether ecdysteroid signaling has a pivotal role in the OA-Oamb-Ca^2+^-dependent GSC increase. As we have previously reported, the RNAi of *neverland* (*nvd*), which encodes an ecdysteroidogenic enzyme (Yoshiyama-Yanagawa et al., 2011; Yoshiyama et al., 2006) in the escort cells, suppressed the mating-induced GSC increase (Figure 3A)(Ameku et al., 2017; Ameku and Niwa, 2016). We also found a similar phenotype in the RNAi of *ecdysone receptor* (*EcR*) in the escort cells (Figure 3A). To assess the requirement of ecdysteroid biosynthesis and signaling in OA-induced GSC increase, we employed an *ex vivo* experiment. Interestingly, the OA-mediated GSC increase was not observed in *nvd* RNAi ovaries (*c587>nvd^RNAi^*) (Figure 3B). Moreover, this impairment of GSC increase in *c587>nvd^RNAi^* was restored with the administration of 20-hydroxyecdysone (20E) in the culture media. On the other hand, 20E treatment without OA did not induce GSC increase in the control ovaries (Figure 3B). This observation is consistent with our previous study showing that wild-type virgin females fed with 20E did not exhibit any increase in GSCs (Ameku and Niwa, 2016). Therefore, 20E is required in the OA-mediated increase in GSCs; however, by itself, 20E is not sufficient to induce GSC increase.

**Figure 3.**
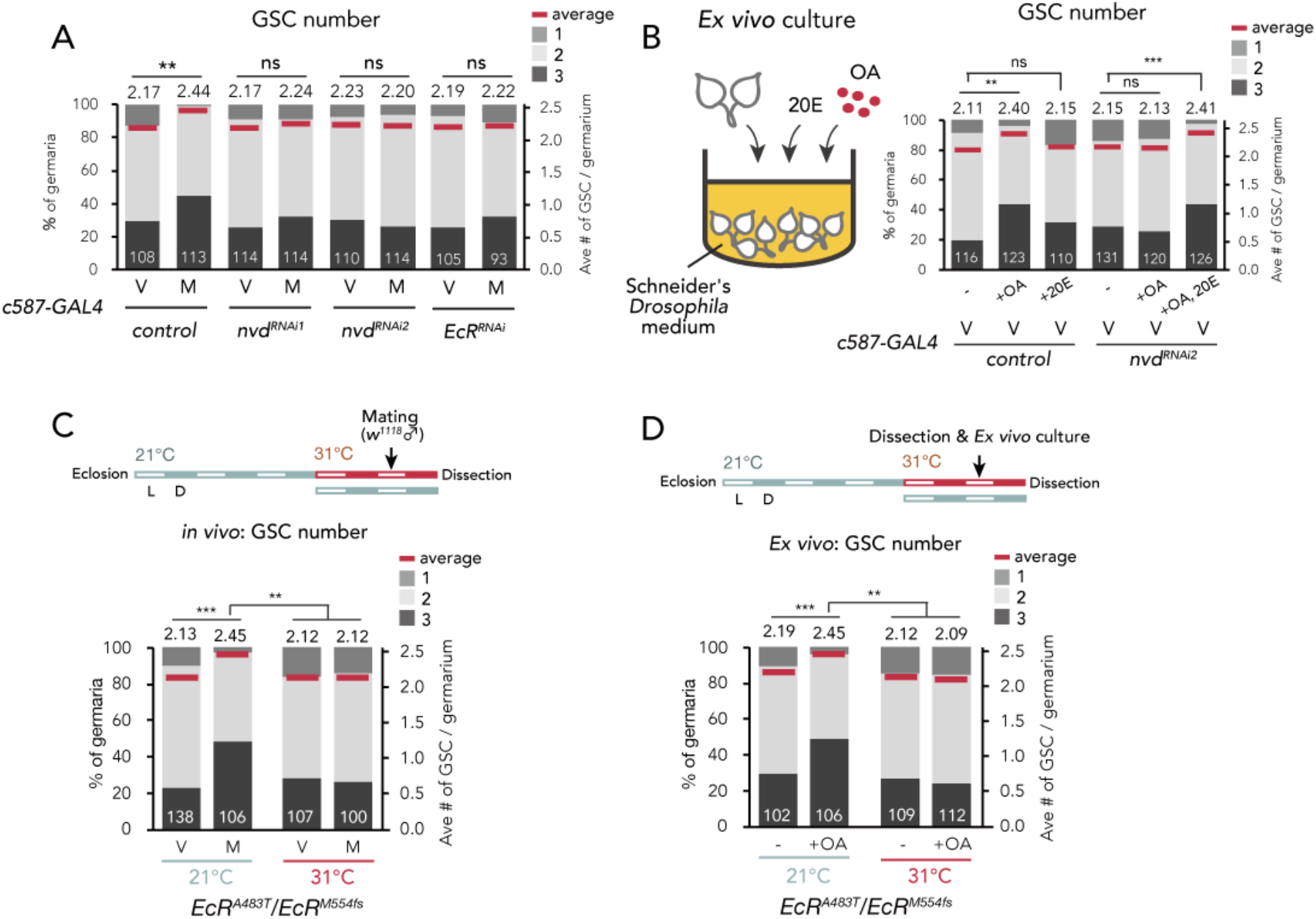
Ecdysteroid signaling is necessary for OA-mediated GSC increase. (A-D) Frequencies of germaria containing 1, 2, and 3 GSCs (left vertical axis) and the average number of GSCs per germarium (right vertical axis) in virgin (V) and mated (M) female flies. *c587>+* flies were used as the control in A and B. The number of germaria analyzed is indicated inside the bars. (A) GSC number of *nvd* and *EcR* RNAi flies *in vivo*. (B) Virgin ovaries were cultured *ex vivo* with or without OA and 20E (+OA, +20E, -), and then the GSC number was determined. (C-D) Experiments using a temperature-sensitive allele *EcR^A483T^*. 21 °C and 31 °C were used as the permissive and restrictive temperatures, respectively. Flies were cultured at 21 °C and transferred to 29 °C 1 d prior to the assays (L; light, D; dark). (C) GSC number *in vivo*. (D) Virgin ovaries were cultured *ex vivo* with or without OA (+OA, -). The number of germaria analyzed is indicated inside the bars. Wilcoxon rank sum test with Holm’s correction was used for statistical analysis. ***P ≤ 0.001 and **P ≤ 0.01; NS, non-significant (P > 0.05).

To assess the role of *EcR* in downstream OA signaling, we utilized the temperature-sensitive allele and the null allele of *EcR* (*EcR^A483T^* and *EcR^M54fs^*, respectively) (Bender et al., 1997). At a restrictive temperature of 31°C, the mating-induced GSC increase was suppressed in *EcR^A483T^*/*EcR^M54fs^* flies, consistent with our previous report (Ameku and Niwa, 2016) (Figure 3C). In the *ex vivo* experiment, the OA-dependent GSC increase was also suppressed in *EcR^A483T^*/*EcR^M54fs^* ovaries (Figure 3D). These results suggest that OA-Oamb-Ca^2+^ signaling requires the ovarian ecdysteroid signaling.

### Matrix metalloprotease 2 acts downstream of OA-Oamb-Ca^2+^ signaling

So far, we have identified four components with indispensable roles in mating-induced GSC increase, namely OA, Oamb, Ca^2+^, and ecdysteroids. Interestingly, recent studies have reported that they are also essential in the follicle rapture in *D. melanogaster* ovary (Deady and Sun, 2015; Knapp and Sun, 2017). In this process, Matrix metalloprotease 2 (Mmp2), a membrane-conjugated protease, acts downstream of the OA-Oamb-Ca^2+^ signaling pathway in mature follicle cells (Deady et al., 2015). Because recent studies have indicated the expression of *Mmp2* in niche cells, including escort cells (Deady et al., 2017, 2015; Wang and Page-McCaw, 2014), we examined Mmp2 function in the escort cells using RNAi of *Mmp2* with *c587-GAL4*. Similar to *c587>Oamb^RNAi^*, *c587>Mmp2^RNAi^* impaired mating-induced GSC increase (Figure 4A). Notably, *Mmp2* knock-down in cap cells by *bab-GAL4* also suppressed GSC increase, suggesting that Mmp2 acts in both escort cells and cap cells to induce GSC increase (Figure 4-figure supplement 1A). We also found that *Mmp2* RNAi in the ovarian-somatic cells by another *GAL4* (*tj>Mmp2^RNAi1^*) resulted in the failure of mating-induced GSC increase, whereas *Mmp2* RNAi in the nervous system (*nSyb>Mmp2^RNAi1^*) and in the follicle cells of stage 14 oocyte (*R44E10>Mmp2^RNAi1^*) had no effect on GSC increase (Figure 4-figure supplement 1B). These results suggest that Mmp2 in the escort cells and cap cells are necessary to induce post-mating GSC increase.

**Figure 4.**
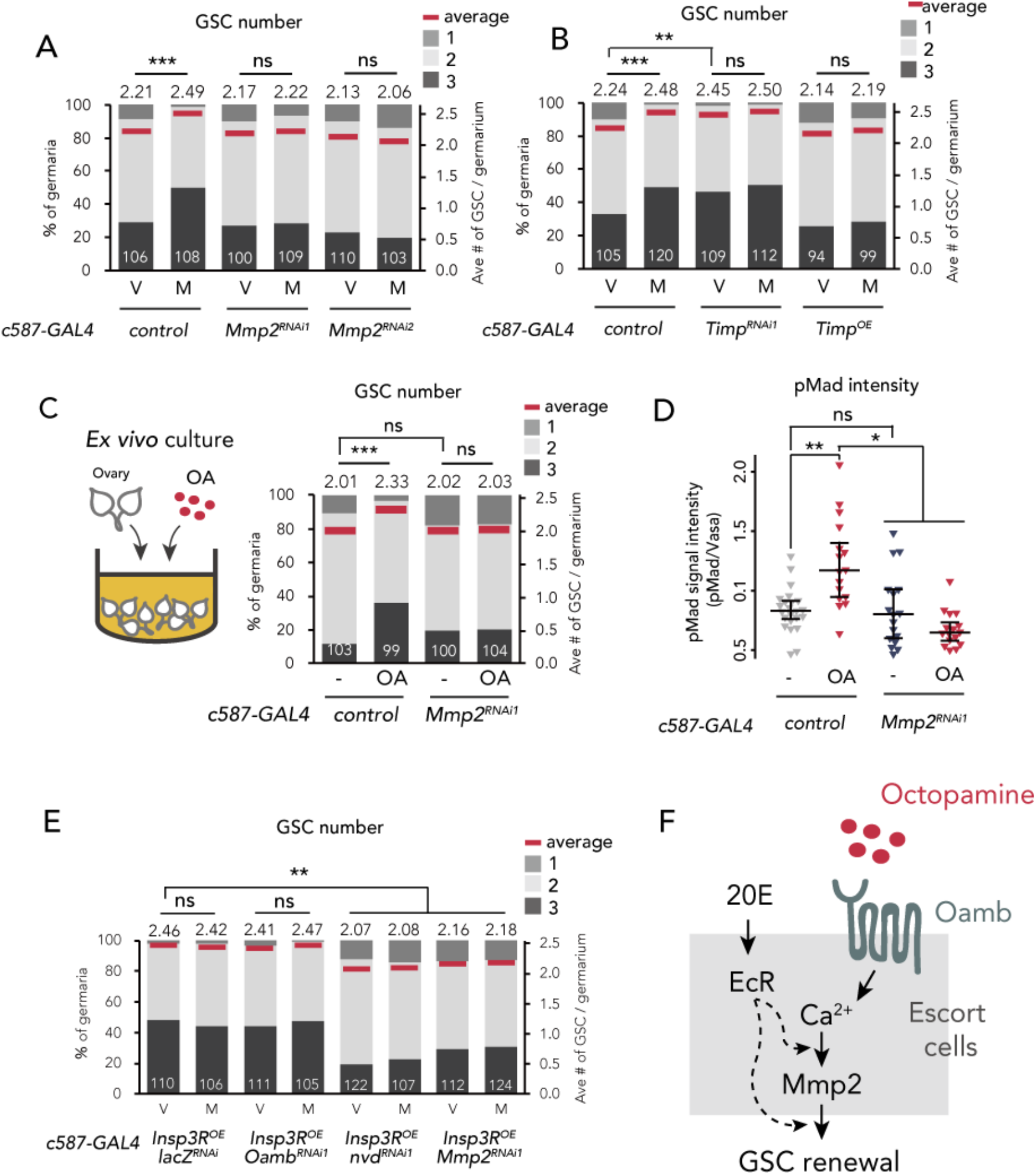
Mmp2 is necessary for OA-mediated GSC increase. (A-C, E) Frequencies of germaria containing 1, 2, and 3 GSCs (left vertical axis) and the average number of GSCs per germarium (right vertical axis) in virgin (V) and mated (M) female flies. *c587>+* flies were used as the control in A, B, C, and D. The number of germaria analyzed is indicated inside the bars. (A) *Mmp2* RNAi by *c587-GAL4* driver. (B) RNAi and the overexpression of *Timp* by *c587-GAL4* driver. (C) *Ex vivo* culture experiment using *c587>Mmp2^RNAi^*. OA was added into the *ex vivo* culture medium. Cultured with or without OA (+OA, -, respectively) is indicated under each bar. (D) Quantification of the relative pMad intensity in GSCs of the *ex vivo* cultured ovaries normalized to pMad intensity in CBs. Cultured with or without OA (+OA, -) is indicated under each bar. For the quantification of pMad intensity, the cell boundaries of GSCs and CBs were determined using anti-Vasa staining (n>15). The 3 horizontal lines for each data sample indicate lower, median, and upper quartiles. (E) *Oamb*, *nvd*, or *Mmp2* RNAi in the genetic background of *c587>Insp3R* overexpression. (F) A model of signaling in the escort cell to induce the mating-induced GSC increase. Oamb in the escort cells receives OA, and induce [Ca^2+^]_i_ in the cells. The [Ca^2+^]_i_ induces GSC increase via Mmp2. Ecdysteroid signaling is also involved in this process. Wilcoxon rank sum test with Holm’s correction was used. ***P ≤ 0.001, **P ≤ 0.01, and *P ≤ 0.05; NS, non-significant (P > 0.05).

In the GSC niche cells in *D. melanogaster*, *Tissue inhibitors of metalloproteases* (*Timp*) gene encoding an endogenous protease inhibitor of Mmp2 is expressed (Gomis-Rüth et al., 1997; Page-McCaw et al., 2003; Pearson et al., 2016). The knock-downn of *Timp* in escort cells induced GSC increase even in virgin females, whereas its overexpression suppressed mating-induced GSC increase (Figure 4B). Consistent with *Mmp2* kncokdown, *Timp* knock-down in cap cells (*bab>Timp^RNAi2^*) increased the GSC number in virgin females, whereas its knock-down in the follicle cells of stage 14 oocyte (*R44E10>Timp^RNAi2^*) had no effect (Figure 4-figure supplement 1C). These data suggest that Mmp2 activity in the GSC niche cells is necessary for mating-induced GSC increase and is suppressed by *Timp* in virgin females.

We next examined whether Mmp2 is necessary for OA-induced GSC increase. Our *ex vivo* culture experiment revealed that OA-induced GSC increase was suppressed in *c587>Mmp2^RNAi^* flies, suggesting that Mmp2 acts downstream of OA signaling (Figure 4C). Moreover, the OA-dependent upregulation of pMad level in GSCs was suppressed in *c587>Mmp2^RNAi^* flies (Figure 4D). Notably, in virgin females, *Mmp2* RNAi affected neither the GSC number nor the cap cell number, indicating that *Mmp2* RNAi does not influence the overall niche architecture (Figure 4 – figure supplement 1D). Mmp2 in the mature follicle cells cleaves and downregulates collagen VI, also known as Viking (Vkg) and is the major component of the basement membrane (Deady et al., 2017; Wang et al., 2008). However, *Mmp2* RNAi had no effect on Vkg::GFP level around the cap cells (Figure 4 – figure supplement 1E). Therefore, the suppression of mating-induced GSC increase in *Mmp2* RNAi does not likely depend on the collagen VI level.

To examine the epistasis of OA/Oamb-Ca^2+^ signaling, ecdysteroid signaling, and Mmp2, we knocked down *Oamb, nvd*, or *Mmp2* in *c587>Insp3R^OE^* genetic background, where Ca^2+^ signaling was forcedly upregulated. Whereas the *Oamb* RNAi did not suppress GSC increase in virgin female (*c587>Insp3R^OE^, Oamb^RNAi^*), the RNAi of *nvd* or *Mmp2* suppressed GSC increase even when Ca^2+^ signaling were activated (*c587>Insp3R^OE^, nvd^RNAi^* or *c587>Insp3R^OE^, Mmp2^RNAi^*) (Figure 4E). These results suggest that ecdysteroid signaling and Mmp2 act downstream of Ca^2+^ signaling in OA-induced GSC increase. Taken together, both the OA-Oamb signaling and the downstream Ca^2+^ signaling regulate the mating-induced GSC increase via Mmp2 and ecdysteroid signaling (Figure 4F).

### Octopamine from *dsx^+^ Tdc2^+^* neurons regulates the mating-induced GSC increase

To examine the *in vivo* role of OA in mating-induced GSC increase, we silenced the expression of *Tyrosine decarboxylase 2* (*Tdc2)* and *Tyramine β hydroxylase* (*TβH)* genes, which code for enzymes responsible for OA biosynthesis (Cole et al., 2005; Monastirioti et al., 1996) (Figure 5A), with *Tdc2-GAL4* driver-mediated RNAi (*Tdc2>Tdc2^RNAi1^, Tdc2> TβH ^RNAi1^*). Similar to the phenotype of *Oamb* RNAi, *Tdc2* or *TβH* RNAi with *Tdc2-GAL4* or *nSyb-GAL4* impaired the mating-induced GSC increase (Figure 5A and Figure 5 – figure supplement 1A-B). Moreover, the impairment of GSC increase was restored when *Tdc2* or *TβH* RNAi flies were fed with food supplemented with OA (Figure 5A), supporting our hypothesis that OA is responsible for the mating-induced GSC increase *in vivo*.

**Figure 5.**
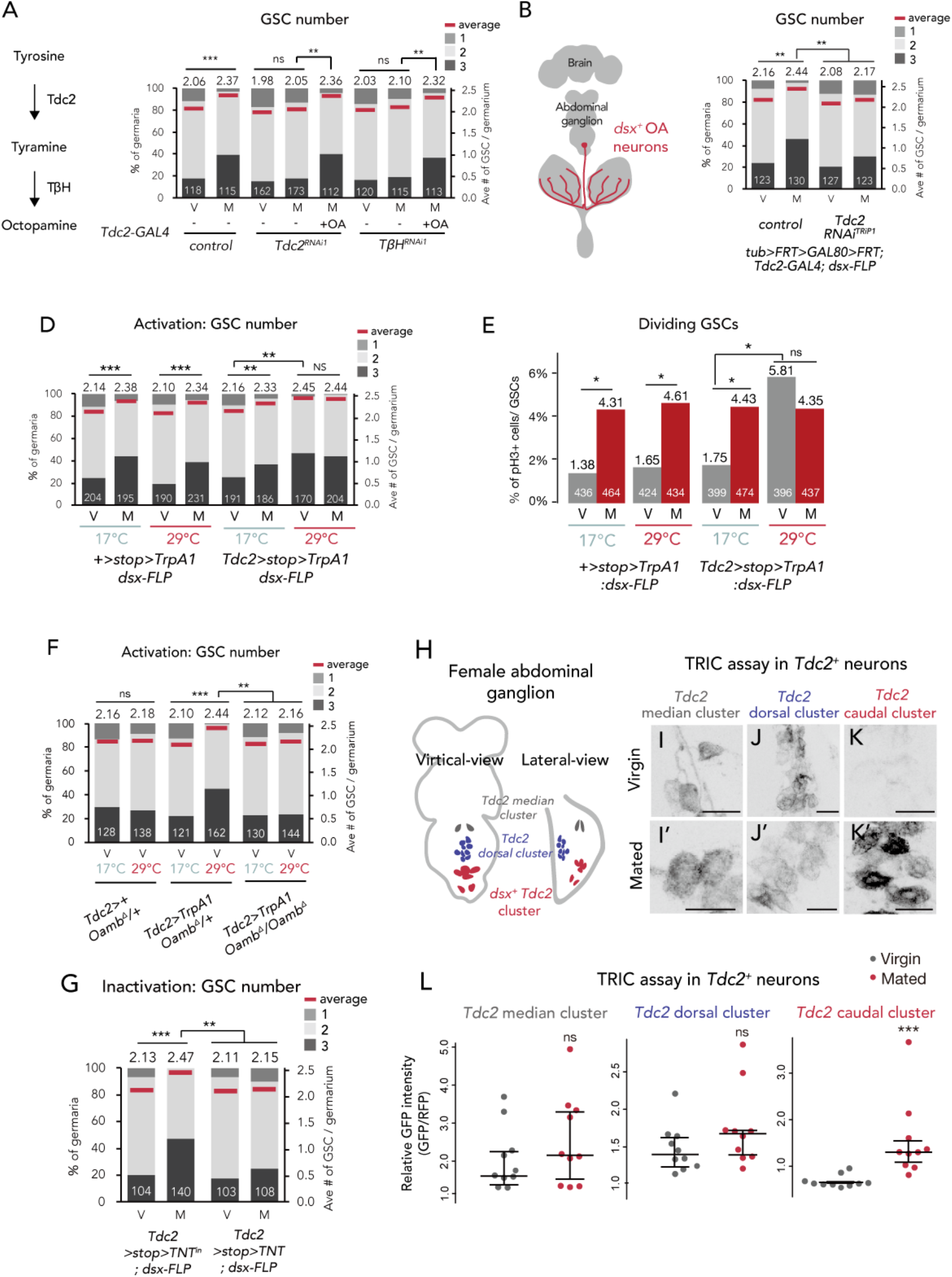
Ovary-projecting OA neurons control the GSC increase. (A, C-D, F-G) Frequencies of germaria containing 1, 2, and 3 GSCs (left vertical axis) and the average number of GSCs per germarium (right vertical axis) in virgin (V) and mated (M) female flies. The number of germaria analyzed is indicated inside the bars. (A) RNAi of *Tdc2* and *TβH* by *Tdc2-GAL4.* OA was added into the standard food. (B) A schematic drawing of *Drosophila* central nervous system and the ovary-projecting OA neurons with the *dsx^+^* OA neurons projecting to the ovary. (C) *Tdc2* RNAi in *dsx^+^ Tdc2^+^* neurons with the genotype indicated. (D-E) *TrpA1*-mediated activation of *dsx^+^ Tdc2^+^* neurons. 17 °C and 29 °C were used as the permissive and restrictive temperatures, respectively, of TrpA1 channel. (D) GSC number. (E) The ratio of pH3^+^ GSCs and total GSCs. (F) The activation of *Tdc2^+^* neurons with *OambΔ* genetic background. (G) The inactivation of *dsx^+^ Tdc2^+^* neurons. (H) Illustration showing the location of 3 clusters of *Tdc2^+^* neurons in the caudal part of the abdominal ganglion (I-K, I’-K’). Negative images of TRIC labelling (anti-GFP) in the abdominal ganglions of virgin (I-K) and mated females (I’-K’) of TRIC (*Tdc2>UAS-mCD8::RFP, UAS-p65AD::CaM LexAop2-mCD8::GFP; nSyb-MKII::nlsLexADBDo;UAS-p65AD::CaM*) flies, indicating intracellular Ca^2+^ transients. Scale bars, 20 µm. (L) The GFP intensities from the *Tdc2^+^* median cluster, *Tdc2^+^* dorsal cluster, and *dsx^+^ Tdc2^+^* cluster of TRIC females show Ca^2+^ activity in virgin (grey) and mated females (red). Wilcoxon rank sum test was used for A, C, D, F, G, and L. Fisher’s exact test with Holm’s correction was used for E. ***P ≤ 0.001, **P ≤ 0.01, and *P ≤ 0.05; NS, non-significant (P > 0.05).

Because *Tdc2-GAL4* is active in the nervous system (Busch et al., 2009; Pauls et al., 2018), we then identified which neurons secrete OA to the escort cells. *D. melanogaster* has more than 70–100 OAergic neurons dispersed throughout the nervous system (Monastirioti, 2003; Schwaerzel et al., 2003). Among them, we were particularly interested in a small subset innervating to the reproductive system (Figure 5B) as several recent studies have revealed that these neurons regulate mating behavior, egg laying, and ovarian muscle contraction (Heifetz et al., 2014; Kurz et al., 2017; Lee et al., 2003; Middleton et al., 2006; Rezával et al., 2014; Rubinstein and Wolfner, 2013). The ovary-projecting OAergic neurons are *doublesex* (*dsx*)*^+^* and *Tdc2^+^* double-positive (Rezával et al., 2014) (Figure 5B). Therefore, to manipulate the gene expression of *dsx^+^ Tdc2^+^* neurons only, we implemented a FLP/FRT intersectional strategy. Using *dsx-FLP* (Rezával et al., 2014), we could detect *GFP* expression only in the *dsx^+^ Tdc2^+^* neurons innervating to the ovary, whose cell bodies are located on a caudal part of the abdominal ganglion (Rezával et al., 2014) (Figure 5-figure supplement 1C-E). We next knocked-down *Tdc2* in *dsx^+^ Tdc2^+^* neurons by RNAi (*tub>GAL80>Tdc2^RNAi^; dsx-FLP*) and found that these RNAi flies failed to increase GSC number after mating (Figure 5C). These data suggest that only a small subset of *dsx^+^ Tdc2^+^* neurons controls the mating-induced GSC increase.

To assess whether the activity of *dsx^+^ Tdc2^+^* neurons affects the GSC number, we overexpressed *TrpA1*, a temperature-sensitive cation channel gene, in the *dsx^+^ Tdc2^+^* neurons only. In *Tdc2>stop>TrpA1;dsx-FLP* flies, we can tightly control the *TrpA1* expression in the *dsx^+^ Tdc2^+^* neurons only (Rezával et al., 2014). Both the control flies and *TrpA1*-overexpressing flies at permissive temperature (17 °C) had the normal GSC number in virgin and mated females. On the other hand, at the restrictive temperature (29 °C), the *TrpA1*-overexpressing flies, even the virgin ones, had more GSCs (Figure 5D). We also found that *Tdc2>stop>TrpA1; dsx-FLP* virgin females at the restrictive temperature had increased GSC frequency in the M phase (Figure 5E). Importantly, the *TrpA1*-mediated activation of *Tdc2* neurons did not induce the GSC increase in loss-of -*Oamb*-function females (*Tdc2>TrpA1; Oamb^Δ^/Oamb^Δ^*) (Figure 5F), suggesting that the *TrpA1*-mediated GSC increase requires Oamb.

Furthermore, we employed the *Tetanus toxin light chain* (*TNT*) to inhibit neuronal activity (Sweeney et al., 1995). When we overexpressed *TNT* in *dsx^+^ Tdc2^+^* neurons only, the mating-induced GSC increase was suppressed in mated females as compared with the control, whose inactivated *TNT^in^* was overexpressed (Figure 5G). Taken together, these findings suggest that the mating-induced GSC increase is mediated by the neuronal activity of *dsx^+^ Tdc2^+^* neurons innervating to the ovary.

### *dsx^+^ Tdc2^+^* neurons are activated after mating

Because the *dsx^+^ Tdc2^+^* neuronal activity has a significant role in mating-induced GSC increase, we next examined whether these neurons change their activity before and after mating. We monitored the neuronal activity using an end-point Ca^2+^ reporting system, the transcriptional reporter of intracellular Ca^2+^ (TRIC) (Gao et al., 2015). TRIC is designed to increase the *GFP* expression in proportion to [Ca^2+^]_i_. We classified female *Tdc2^+^* neurons in the caudal part of the abdominal ganglion into 3 clusters based on their location and morphology. We designated the 3 clusters of these *Tdc2^+^* neurons as the *Tdc2^+^* median, *Tdc2^+^* dorsal, and *Tdc2^+^* caudal clusters (Figure 5H). Among them, the position of first 2 clusters are not similar to that of *dsx^+^ Tdc2^+^* neurons, whereas that of the *Tdc2^+^* caudal cluster is similar (Rezával et al., 2014). In virgin females, we detected robust TRIC signals in the *Tdc2^+^* median and *Tdc2^+^* dorsal clusters but not in the *Tdc2^+^* caudal cluster (Figure 5I-L). In contrast, 24 h after mating, we observed a significant increase in the TRIC signal in the *Tdc2^+^* caudal cluster, whereas those in the *Tdc2^+^* median and *Tdc2^+^* dorsal clusters were not changed in virgin and mated females (Figure 5I-L). This result suggests that the *Tdc2^+^* caudal cluster, which are likely *dsx^+^ Tdc2^+^* neurons, is significantly activated after mating.

### The activity switch of *dsx^+^ Tdc2^+^* neurons are regulated by the Sex peptide sensory neurons via acetylcholine signaling

Our previous study revealed that the mating-induced GSC increase is mediated by the male seminal fluid protein SP (Ameku and Niwa, 2016). SP is received by SPR in a small number of SPSNs, followed by a neural silencing of SPSNs (Häsemeyer et al., 2009; Yapici et al., 2008). Notably, SPSNs project their arbors into a caudal part of the abdominal ganglion, where the cell bodies of the *dsx^+^ Tdc2^+^* cluster neurons are located (Rezával et al., 2014, 2012). Therefore, we examined whether SPSNs physically interact with *Tdc2^+^* neurons in the abdominal ganglion by performing the GFP Reconstitution Across Synaptic Partners (GRASP) analysis (Feinberg et al., 2008; Gordon and Scott, 2009), in which two complementary fragments of *GFP* were expressed in SPSNs and *Tdc2^+^* neurons. GRASP signals were detected in the abdominal ganglion (Figure 6A), suggesting that the axon termini of SPSNs and the cell bodies and/or dendrites of *Tdc2^+^* neurons contact each other likely through synaptic connections.

**Figure 6.**
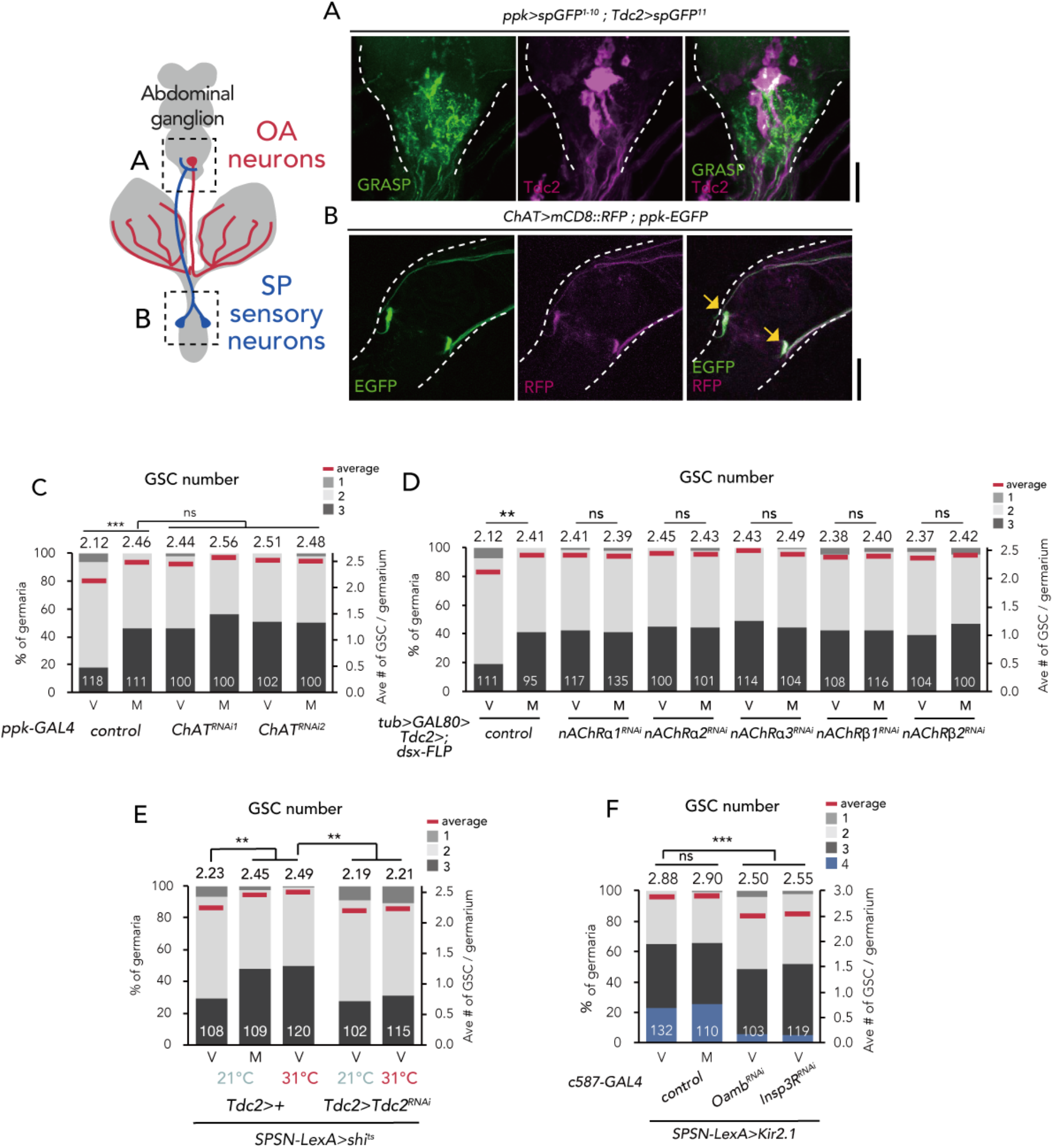
SPSNs control GSC increase through OA neurons. (A) Neuronal proximity of SPSNs and *Tdc2^+^* neurons in the abdominal ganglion of female flies stained with anti-Tdc2 (magenta). Note that reconstituted GFP (GRASP) signal was detected in the caudal part of the abdominal ganglion surrounded by broken white lines. Scale bar, 25 µm. (B) Cell bodies of SPSNs (yellow arrows) of *ChAT-GAL4; UAS-mCD8::RFP*; *ppk-EGFP* virgin females. Note that mCD8::RFP and EGFP signals overlapped in the cell bodies of SPSNs. White broken lines outline the oviduct. Scale bar, 25 µm. (C-F) Frequencies of germaria containing 1, 2, 3 and 4 GSCs (left vertical axis) and the average number of GSCs per germarium (right vertical axis) in virgin (V) and mated (M) female flies. The number of germaria analyzed is indicated inside the bars. (C) *ChAT* RNAi by *ppk-GAL4*. (D) RNAi of *nAChR*s in *dsx^+^ Tdc2^+^* neurons. (E) RNAi of *Tdc2* by *Tdc2-GAL4* along with the silencing of SPSNs. 21 °C and 31 °C were used as the permissive and restrictive temperatures, respectively, of *shibire^ts^ (shi^ts^)*. (F) RNAi of *Oamb* and *Insp3R* by *c587-GAL4* along with the silencing of SPSNs. *Kir2.1* was used in this experiment. Note that frequencies of germaria containing 4 GSCs increased in these genotypes. Wilcoxon rank sum test with Holm’s correction was used for C, D, E and F. ***P ≤ 0.001 and **P ≤ 0.01; NS, non-significant (P > 0.05).

Because SPSNs have been implied as cholinergic neurons (Rezával et al., 2012), we next examined the expression of *Choline acetyltransferase* (*ChAT*)*-GAL4* in SPSNs. *ChAT* encodes an acetylcholine biogenic enzyme (Greenspan, 1980). The SPSNs located on the oviduct, which also co-express *pickpocket* (*ppk*) and *fruitless* (*fru*), are particularly crucial for inducing the major behavioral changes in female flies after mating (Ameku and Niwa, 2016; Rezával et al., 2012). By using *UAS-mCD8::RFP* with *ChAT -GAL4* alongside *ppk-EGFP,* we confirmed that the *ppk-EGFP–*positive population near the oviduct were co-labeled by mCD8::RFP (Figure 6B), consistent with the speculation that SPSNs are cholinergic.

We next counted the GSC number in *ChAT* RNAi flies using *ppk-GAL4. ppk>ChAT ^RNAi^* virgin flies had more GSCs compared with the control (Figure 6C). In addition, mating did not induce GSC increase in *ppk>ChAT^RNAi^* files, suggesting that the acetylcholine released from SPSNs is responsible for suppressing the GSC increase.

To further ascertain whether the acetylcholine released from SPSNs is received by *dsx^+^ Tdc2^+^* neurons to mediate mating-induced GSC increase, we focused on the fast-ionotropic nicotinic acetylcholine receptors (nAChR), which belong to the Cys-loop receptor subfamily of ligand-gated ion channels (Breer and Sattelle, 1987; Gundelfinger and Hess, 1992; Lee and O’Dowd, 1999). In *D. melanogaster,* 10 genes coding *nAChR* subunits (*nAChR*s) have been identified. Among these, we focused on *nAChRα1*, *nAChRα2, nAChRα3, nAChRβ1* and *nAChRβ2* because the knock-down of these genes in *dsx^+^ Tdc2^+^* (*tub>GAL80>, dsx-FLP; Tdc2-GAL4*) or *Tdc2* neurons (*Tdc2-GAL4*) increased the GSC number in virgin females similar to *ppk>ChAT^RNAi^* (Figure 6D and Figure 6-figure supplement 1A). We then confirmed the expression of these *acetylcholine receptor* genes in *Tdc2* neurons by generating a knock-in *T2A-GAL4* line as previously described (Kondo et al., 2020) for each 5 *nAChR*s and observed their expression with *UAS-mCD8::GFP*. All of the five *knock-in-GAL4* expression were detected in anti-Tdc2 positive neurons around the ovary, suggesting that the ovary projecting *dsx^+^ Tdc2^+^* neurons expresses these *nAChR*s (Figure 6-figure supplement 1B-F).

To confirm the role of nAChR in mating-induced GSC increase, we generated *nAChRα1* complete loss-of-function genetic allele by CRISPR/Cas9 technology (Kondo and Ueda, 2013) (Figure 6-figure supplement 2A). Similar to *nAChRα1* RNAi females, the *nAChRα1* transheterozygous mutant virgin females (*nAChRα1^228^/nAChRα^326^*) had more GSCs compared with the controls (Figure 6-figure supplement 2B). In addition, the GSC increase of *nAChRα1^228^/ nAChRα^326^* was restored by the overexpression of *nAChRa1* in *Tdc2^+^* neurons (*Tdc2>nAChRα1; nAChRα1^228^/nAChRα^326^*) (Figure 6-figure supplement 2C). These data support our hypothesis that acetylcholine signaling in *Tdc2* neurons has a negative role in mating-induced GSC increase.

We next assessed relationship between SPSNs, *Tdc2^+^* neurons, and OA-Oamb-Ca^2+^ signaling in ovarian cells. The silencing of SPSNs neuronal activity (*SPSNs-LexA*>*LexAop-shi^ts^*) increased the GSC number in virgin females (Figure 6E). Upon SPSNs silencing, *Tdc2* RNAi by *Tdc2-GAL4* reduced the GSC number (Figure 6E), suggesting that *Tdc2^+^* neurons act downstream of SPSNs. In addition, the GSC increase through the silencing of SPSNs (*SPSNs-LexA>LexAop-Kir2.1*) was suppressed by *Oamb* or *Insp3R* RNAi in the escort cells (Figure 6F), suggesting that OA-Oamb-Ca^2+^ signaling in ovarian cells acts downstream of SPSNs. Overall, our findings revealed the novel neuronal relay in response to mating regulating the female GSC increase in the ovary before/after mating (Figure 7).

**Figure 7.**
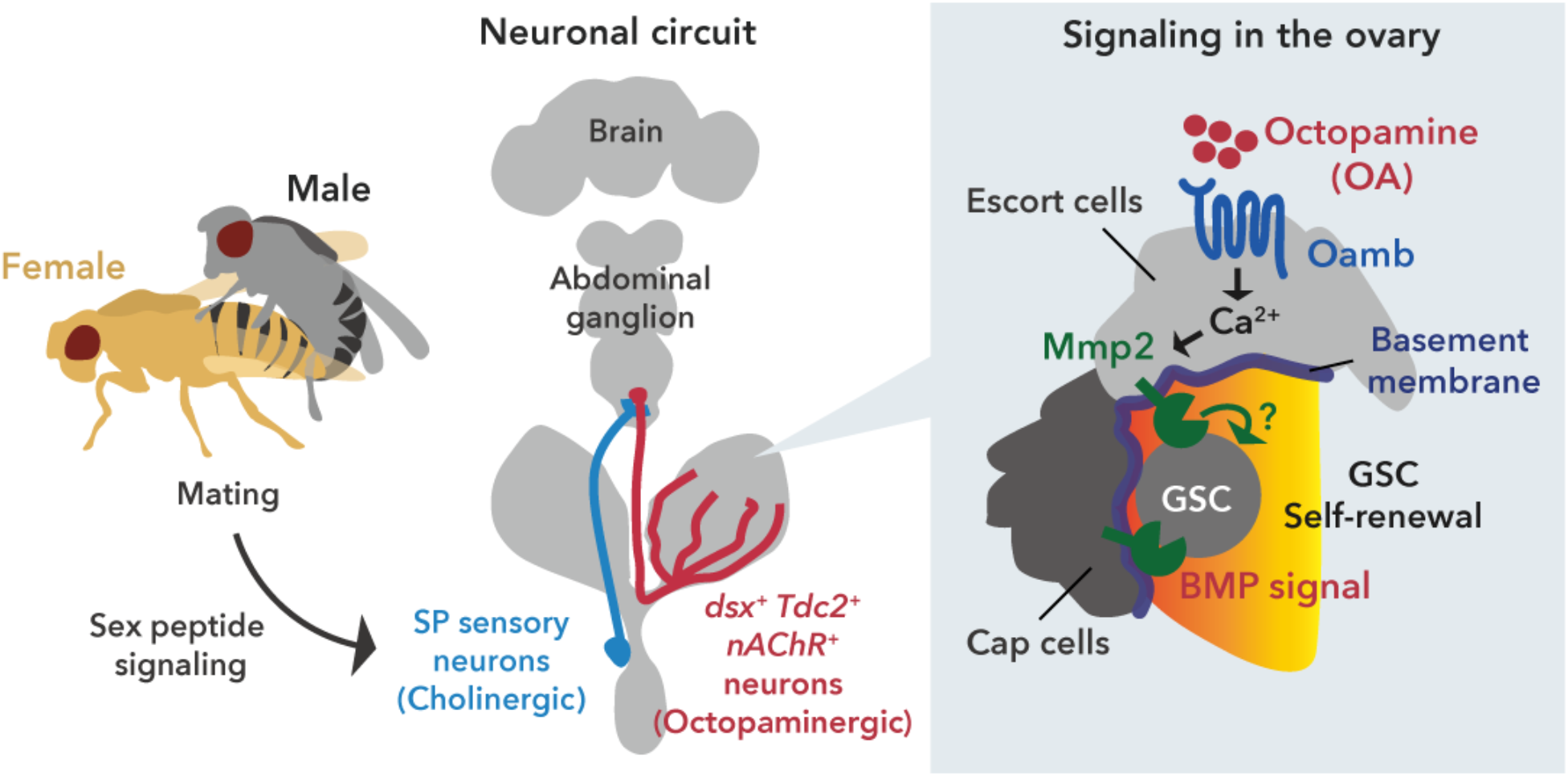
Neuronal octopamine signaling, followed by Oamb-Ca^2+^-Mmp2 signaling, regulates the mating-induced GSC proliferation. The illustration is the proposed working model from our findings here. SP signaling and SP sensory neurons activate *dsx^+^ Tdc2* neurons via acetylcholine signaling. The octopamine released from *dsx^+^ Tdc2* neurons is received by the Oamb in escort cells and then activates intracellular Ca^2+^ flux. The OA-mediated signaling increases pMad levels in GSCs to evoke mating-induced GSC increase via Mmp2.

## Discussion

In this study, we report that the mating-induced GSC proliferation in female *D. melanogaster* is regulated by OAergic neurons directly projecting to the ovary. From our *in vivo* and *ex vivo* experiments, we propose the following model to explain the mating-induced GSC proliferation. After mating, the male seminal fluid SP is transferred into the female uterus, stimulating SPR-positive neurons. As the liganded SPR silences the neuronal activity of SPSNs (Häsemeyer et al., 2009), the acetylcholine released from SPSNs is suppressed. As SPSNs and *dsx^+^ Tdc2^+^* neurons are directly connected, this suppression directly modulates *dsx^+^ Tdc2^+^* neuronal activity. Because we have showed that nAChRs in *dsx^+^ Tdc2^+^* neurons exhibit an inhibitory effect with an unknown mechanism (to be discussed later), the inactivation of nAChRs in the absence of acetylcholine results in the activation of *dsx^+^ Tdc2^+^* neurons in mated females. As a consequence, OA is released from *dsx^+^ Tdc2^+^* neurons, received by Oamb, induces [Ca^2+^]_i_ in the escort cells, and finally activates the Mmp2 enzymatic activity. The activity of Mmp2 positively regulates the Dpp-mediated niche signaling, thereby leading to mating-induced GSC proliferation (Figure 7).

### The ovary-projecting neuron-dependent GSC control

We have showed that the activation of the ovary-projecting *dsx^+^ Tdc2^+^* neurons is necessary and sufficient to induce GSC increase. According to our proposed model, the OA from *dsx^+^ Tdc2^+^* neurons is directly received by the escort cells in the germarium. However, from an anatomical point of view, the *dsx^+^ Tdc2^+^* neurons project to the distal half of the ovary but not to the germarium (Figure 5-figure supplement 1E). This disagreement can be attributed to the characteristic volume transmission of monoamine neurotransmitters. In other words, neurotransmitters act at a distance well beyond their release sites from cells or synapses (Fuxe et al., 2010). Therefore, the OA secreted from the terminals of *dsx^+^ Tdc2^+^* neurons can reach the germarium located at the most proximal part of the ovary. We also emphasize that *Oamb* is expressed in the escort cells in the germarium, and OA treatment evokes [Ca^2+^]_i_ elevation in an Oamb-dependent manner.

Several previous studies have revealed that OA signaling has a pivotal role in the reproductive tissues other than germarium, such as mature follicle cells, oviduct, and ovarian muscle, to promote ovulation, oviduct remodeling, and ovarian muscle contraction, respectively (Deady and Sun, 2015; Heifetz et al., 2014; Lee et al., 2009; Middleton et al., 2006; Rezával et al., 2014). Therefore, it is likely that the *dsx^+^ Tdc2^+^* neurons orchestrate multiple different events during oogenesis in response to mating stimulus. Because a mated female needs to activate oogenesis to continuously produce eggs in concert with sperm availability, it is reasonable that the ovary-projecting neurons switch on the activity of the entire process of reproduction.

### Mmp2 activity in the GSC niche for GSC control

Based on our present study and several previous studies (Deady et al., 2015; Deady and Sun, 2015; Knapp and Sun, 2017), the OA-Oamb-Ca^2+^-Mmp2 axis is required for GSC increase and follicle rupture, both of which are induced by mating stimuli in *D. melanogaster*. In both cases, Mmp2 enzymatic activity is likely to be essential, as the overexpression of *Timp* encoding a protein inhibitor of Mmp2 suppresses GSC increase, as well as follicle rupture. Mmp2 in mature follicle cells cleaves and downregulates Viking/collagen VI (Deady et al., 2017; Wang et al., 2008). In fact, several previous studies have revealed that Viking/collagen VI is required for GSC maintenance in female *D. melanogaster* (Van De Bor et al., 2015; Wang et al., 2008). However, we observed no significant change in Viking/Collagen VI levels in the germarium between the control and *Mmp2* RNAi flies (Figure 4-figure supplement 1E). Therefore, we concluded that Viking/collagen VI is not a substrate of Mmp2 in the regulation of mating-induced GSC increase. Besides Viking/Collagen VI, Dally-like (Dlp) is another basement membrane protein associated with extracellular matrix and known as the Mmp2 substrate (Wang and Page-McCaw, 2014). Interestingly, *dlp* is expressed in the escort cells (Wang and Page-McCaw, 2014). Moreover, Dlp controls the distribution of Dpp and Wnts, both of which significantly affect GSC self-renewal and differentiation (Wang et al., 2015; Xie and Spradling, 1998). Future research should decipher the exact substrate by which Mmp2 controls Dpp and/or Wnts to modulate GSC behavior in response to mating stimulus.

Another remaining question to be addressed is how Mmp2 enzymatic activity is regulated in oogenesis. Ecdysteroid biosynthesis and signaling in the ovary are necessary but not sufficient for the OA-Oamb-Ca^2+^–mediated GSC increase and follicle rapture (Ameku and Niwa, 2016; Knapp and Sun, 2017). We found that in the regulation of mating-induced GSC increase, ecdysteroid signaling acts downstream of Ca^2+^ signaling (Figure 4F). On the other hand, in the follicle rapture process, ecdysteroid signaling either acts downstream, upstream, or both, of Ca^2+^ signaling. Further, the precise action of ecdysteroid has yet to be elucidated (Knapp and Sun, 2017). The Mmp2-GFP fusion protein level in the mature follicle cells is not changed in the loss-of-*Ecdysone receptor*-function flies, implying that ecdysteroid signaling might regulate Mmp2 enzymatic activity by an unknown mechanism (Knapp and Sun, 2017). Considering the involvement of both the OA-Oamb-Ca^2+^-Mmp2 axis and ecdysteroid biosynthesis, it is very likely that the Mmp2 enzymatic activity is also regulated by the same, unknown mechanism not only in the mature follicle cells to control follicle rapture, but also in the germarium to control mating-induced GSC increase.

### SPSNs-mediated suppression of *dsx^+^/Tdc2^+^* neurons

In many animals, reproduction involves significant behavioral and physiological shifts in response to mating. In female *D. melanogaster*, several post-mating responses are coordinated by SPSNs and its downstream afferent neuronal circuit (Wang et al., 2020), including Stato-Acoustic Ganglion neurons, the ventral abdominal lateral Myoinhibitory peptide neurons, and the efferent *dsx^+^ Tdc2^+^* neurons (Feng et al., 2014; Häsemeyer et al., 2009; Jang et al., 2017; Rezával et al., 2014). Our GRASP analysis indicates a direct synaptic connection between cholinergic SPSNs and OAergic neurons. Moreover, we demonstrated that nAChRs in *dsx^+^ Tdc2^+^* neurons are responsible for the suppression of their neuronal activity in virgin females. However, nAChRs are the cation channels leading to depolarization upon acetylcholine binding, and therefore usually activate neurons (Corringer et al., 2000; Lee and O’Dowd, 1999; Perry et al., 2012). How is the opposite role of nAChRs in *dsx^+^ Tdc2^+^* neuronal activity achieved? One possibility is that acetylcholine-nAChR signaling does not evoke a simple depolarization but rather generates a virgin-specific temporal spike pattern in *dsx^+^ Tdc2^+^* neurons. Interestingly, recent studies demonstrated that the pattern, instead of the frequency, of neuronal firing is significant in adjusting the neuronal activity of clock neurons in *D. melanogaster* (Tabuchi et al., 2018). The firing pattern relies on control of ionic flux by the modulation of Ca^2+^-activated potassium channel and Na^+^/K^+^ ATPase activity. Because whether mating changes the firing pattern of *dsx^+^ Tdc2^+^* neurons remains to be examined, the neuronal activity in SPSNs and the *dsx^+^ Tdc2^+^* neuronal circuit between virgin and mated females are future research areas.

### Interorgan communication among multiple organs to regulate the increase and maintenance of female GSCs

In the last decades, there is growing evidence that GSCs and their niche are influenced by multiple humoral factors (Drummond-Barbosa, 2019; Yoshinari et al., 2019). Based on the data from our current study and previous studies, there are at least 4 crucial humoral factors for regulating the increase and/or maintenance of *D. melanogaster* female GSCs, including DILPs (Hsu et al., 2008; Hsu and Drummond-Barbosa, 2009; LaFever and Drummond-Barbosa, 2005), ecdysteroids (Ables and Drummond-Barbosa, 2010; Ameku et al., 2017; Ameku and Niwa, 2016; König et al., 2011), Neuropeptide F (NPF) (Ameku et al., 2018), and OA (this study). Notably, all of these come from different sources: DILPs are from the insulin-producing cells located in the pars intercerebralis of the central brain; ecdysteroids from the ovary; NPF from the midgut; and OA from the neurons located in the abdominal ganglion. In addition to these identified humoral factors, recent studies also imply that adiponectin and unknown adipocyte-derived factor(s) are essential for GSC maintenance (Armstrong and Drummond-Barbosa, 2018; Laws et al., 2015; Matsuoka et al., 2017). These data clearly indicate that *D. melanogaster* female GSCs are systemically regulated by interorgan communication involving multiple organs. The additional interorgan communication mechanisms that ensure the faithful coupling of the increase and maintenance of GSC to the organism’s external and physiological environments are essential to be investigated in future studies.

To modulate the increase and maintenance of GSC, ecdysteroids are received by both GSCs and niche cells (Ables and Drummond-Barbosa, 2010; König et al., 2011), whereas DILPs, NPF, and OA are received by niche cells. A major signal transduction mechanism of each of these humoral factors have been well characterized, namely phosphoinositide 3-kinase pathway for DILPs-InR signaling, EcR/Ultraspiracle-mediated pathway for ecdysteroid signaling, cAMP pathway for NPF-NPFR signaling (Garczynski et al., 2002), and Ca^2+^ pathway for OA-Oamb signaling. However, it remains unclear whether and how each of these signaling pathways control the production and secretion of the niche signal, as well as its distribution and transduction. In addition, it is important to understand whether and how the multiple system signals are integrated to control the mating-induced increase and maintenance of GSCs.

### Evolutionarily conservation of monoamine-steroid hormone axis to control female reproduction

In *D. melanogaster*, ecdysteroid signaling is essential for the proliferation and maintenance of GSCs and neural stem cells (Ables and Drummond-Barbosa, 2010; Homem et al., 2014; König et al., 2011). In this study, we have identified the ovary-projecting OAergic neurons as new regulators of stem cell homeostasis. Both steroid hormones and OA-like monoamines, such as noradrenaline, are also involved in stem cell regulation in mammals. For example, the mammalian steroid hormone, estrogen, is important in regulating cell division and/or maintenance of hematopoietic stem cells, mammary stem cell, neural stem cells, and hematopoietic stem cells (Asselin-Labat et al., 2010; Bramble et al., 2019; Kim et al., 2016; Nakada et al., 2014). Moreover, noradrenergic neurons, which directly project to the bone marrow, regulate the remodeling of hematopoietic stem cells niche (Ho et al., 2019; Méndez-Ferrer et al., 2010, 2008). Therefore, the steroid hormone- and noradrenergic nerve-dependent control of stem cell homeostasis are likely conserved across animal species. In this regard, *D. melanogaster* reproductive system will further serve as a powerful model to unravel the conserved systemic and neuronal regulatory mechanisms for stem cell homeostasis in animals.

## Materials and methods

### Drosophila strains

Flies were raised on cornmeal-yeast-agar medium at 25 °C. *EcR^A483T^*, temperature-sensitive mutants, were cultured at 29 °C for 1 day prior to the assays. *w^1118^* was used as the control strain.

The genetic mutant stocks used were *EcR^A483T^* (Bloomington Drosophila Stock Center [BDSC] #5799) and *EcR^M554fs^* (BDSC #4894). The protein-trap GFP line of Vkg (Vkg::GFP) was obtained from Kyoto Stock Center (DGRC #110692).

The following *GAL4* and *LexA* strains were used: *c587-GAL4* (Manseau et al., 1997) (gift from Hiroko Sano, Kurume University, Japan), *R44E10-GAL4* (Deady and Sun, 2015) (a gift from Jianjun Sun, University of Connecticut, USA) *, RS-GAL4* (Lee et al., 2009) (a gift from Kyung-An Han, Pennsylvania State University, USA) *, Tk-gut-GAL4* (Song et al., 2014) (a gift from Masayuki Miura, The University of Tokyo, Japan), *nSyb-GAL4* (BDSC #51941), *nSyb-GAL80* (Harris et al., 2015) (a gift from James W. Truman, Janelia Research Campus, USA), *tj-GAL4* (DGRC #104055), *bab-GAL4* (Bolívar et al., 2006) (a gift from Satoru Kobayashi, University of Tsukuba, Japan), *nos-GAL4* (DGRC #107748), *tub>FRT>GAL80>FRT* (BDSC #38879), *Oamb-T2A-GAL4, nAChRα1-T2A-GAL4, nAChRα2-T2A-GAL4, nAChRα3-T2A-GAL4, nAChRβ1-T2A-GAL4, nAChRβ2-T2A-GAL4* (Kondo et al., 2020), *ChAT -GAL4* (BDSC #6793), *ppk-GAL4* (Grueber et al., 2007) (a gift from Hiroko Sano, Kurume University, Japan), and *SPSNs-LexA* (Feng et al., 2014) (a gift from Yong-Joon Kim, Gwangju Institute of Science and Technology, South Korea).

The following *UAS* and *LexAop* strains were used: *20xUAS-6xGFP* (BDSC #52261), *UAS-GFP;UAS-mCD8::GFP* (Ito et al., 1997; Lee and Luo, 1999) (a gift from Kei Ito, University of Cologne, Germany), *UAS-mCD8::RFP* (BDSC #32219), *UAS-CsChrimson* (BDSC #55134), *UAS-Insp3R* (BDSC #30742), *UAS-Timp* (BDSC #58708) (a gift from Andrea Page-McCaw, Vanderbilt University, USA), *UAS>stop>dTrpA1^mcherry^*, *UAS>stop>TNT, UAS>stop>TNT^in^* (von Philipsborn et al., 2011; Yu et al., 2010), *dsx-FLP* (Rezával et al., 2014) (a gift from Daisuke Yamamoto, Advanced ICT Research Institute, National Institute of Information and Communications Technology, Japan) TRiC; *UAS-mCD8::RFP, LexAop2-mCD8::GFP;nSyb-MKII::nlsLexADBDo;UAS-p65AD::CaM* (BDSC:61679), *ppk-eGFP* (Grueber et al., 2003) (a gift from Tadashi Uemura, Kyoto University, Japan), and *LexAop-Kir2.1* (Feng et al., 2014) (a gift from Yong-joon Kim, GIST, Korea).

The RNAi transgenic lines used were as follows: *UAS-LacZ^RNAi^* (a gift from Masayuki Miura, The University of Tokyo, Japan), *UAS-Oamb^RNAi1^*(BDSC #31171), *UAS-Oamb^RNAi2^*(BDSC #31233), *UAS-Oamb^RNAi3^* (Vienna Drosophila Resource Center [VDRC] #106511), *UAS-Octβ1R^RNAi^*(VDRC #110537), *UAS-Octβ2R^RNAi^*(VDRC #104524), *UAS-Octβ3R^RNAi^* (VDRC #101189), *UAS-Insp3R^RNAi^* (BDSC #25937), *UAS-EcR^RNAi^* (VDRC #37059), *UAS-Mmp2^RNAi1^* (BDSC #31371), *UAS-Mmp2^RNAi2^* (VDRC #330303), *UAS-Timp^RNAi1^* (BDSC #61294), *UAS-Timp^RNAi2^* (VDRC #109427), *UAS-Tdc2^RNAi1^* (VDRC #330541), *UAS-Tdc2^RNAi2^* (BDSC #25871), *UAS-Tβh^RNAi1^* (VDRC #107070), *UAS-Tβh^RN1i2^* (BDSC #67968), *UAS-ChAT^RNAi1^* (VDRC #330291), *UAS-ChAT^RNAi2^* (BDSC #25856), *UAS-nAChRα1^RNAi^* (VDRC #48159), *UAS-nAChRα2^RNAi^* (VDRC #101760), *UAS-nAChRα3^RNAi^* (VDRC #101806), *UAS-nAChRβ1^RNAi^* (VDRC #106570), *UAS-nAChRβ2^RNAi^* (VDRC #109450), *UAS-nvd^RNAi1^*, and *UAS-nvd^RNAi2^* (Yoshiyama et al., 2006).

### Generation of *Oamb* and *nAChRα1* genetic loss-of-function mutant strains

The mutant alleles *Oamb^Δ^* (Figure 1-figure supplement 1D), *nAChRα1^228^*, and *nAChRα1^326^* (Figure 6-figure supplement 2A) were created in a *white* (*w*) background using CRISPR/Cas9 as previously described (Kondo and Ueda, 2013). The following guide RNA (gRNA) sequences were used: *Oamb*, 5ʹ-GATGAACTCGAGTACGGCCA-3ʹ, and 5ʹ-GCGATCTCTGGTGCCGCATT-3ʹ; *nAChRα1^228^*, 5ʹ-GGACATCATGCGTGTGCCGG-3ʹ ; *nAChRα1^326^*, 5ʹ-GGGCAGGTAGAAGACCAGAA-3ʹ. The breakpoint detail of *Oamb^Δ^* is described in Figure 1-figure supplement 1D, whereas those of *nAChRα1^228^* and *nAChRα1^326^* are described in Figure 6-figure supplement 2A.

### Generation of *UAS-nAChRα1* transgenic line

The pcDNA3.1 plasmid containing the wild-type *D. melanogaster nAChRα1* coding sequences (*nAChRα1*-pcDNA3.1) was synthesized previously described (Ihara et al., 2018). Briefly, *nAChRα1*-pcDNA3.1 was digested with *EcoR*I and *Not*I, and then the digested *nAChRα1* fragment was ligated with a *EcoR*I-*Not*I–digested pWALIUM10-moe plasmid (Perkins et al., 2015). Transformants were generated using the phiC31 integrase system in the *P{CaryP}attP40* strain (Groth et al., 2004). The *w^+^* transformants of pWALIUM10-moe were established using standard protocols.

### Behavioral assays

Flies were reared at 25 °C and aged for 5–6 d. Virgin female flies were mated overnight to *w^1118^* male flies at 25 °C (10 males and 5–8 females per vial). For the thermal activation assays, flies were first reared at 17 °C for 6 d and transferred to 29 °C. In the case of *EcR^A483T^* mutant assays, flies were transferred to 31 °C for 24 h before mating or *ex vivo* culture.

For OA feeding, newly eclosed virgin females were aged for 4 d in vials with standard food containing 7.5 mg/mL of OA.

### Immunohistochemistry

Tissues were dissected in phosphor buffer serine (PBS) and fixed in 4% paraformaldehyde in PBS for 30 to 60 min at room temperature (RT). The fixed samples were washed 3 times in PBS supplemented with 0.2% Triton X-100, blocked in blocking solution (PBS with 0.3% Triton X-100 and 0.2% bovine serum albumin [BSA]) for 1 h at RT, and incubated with a primary antibody in the blocking solution at 4 °C overnight. The primary antibodies used were chicken anti-GFP (Abcam #ab13970; 1:4,000), rabbit anti-RFP (Medical & Biological Laboratories PM005; 1:2,000), mouse anti-Hts 1B1 (Developmental Studies Hybridoma Bank [DSHB]; 1:50), rat anti-DE-cadherin DCAD2 (DSHB; 1:50), rabbit anti-pH3 (Merck Millipore #06-570; 1:1000), rabbit monoclonal anti-pMad (Abcam #ab52903; 1:1000), mouse anti-Lamin-C LC28.26 (DSHB; 1:10), rabbit cleaved Dcp-1 (Cell Signaling Technology #9578; 1:100), rat anti-Vasa (DSHB; 1:50), rabbit anti-Tdc2 (Abcam #ab128225; 1:2000), Alexa Fluor 546 phalloidin (Thermo Fisher Scientific #A22283; 1:200), and Alexa Fluor 633 phalloidin (Thermo Fisher Scientific #A22284; 1:200). After washing, fluorophore (Alexa Fluor 488, 546 or 633)-conjugated secondary antibodies (Thermo Fisher Scientific) were used at a 1:200 dilution, and the samples were incubated for 2 h at RT in the blocking solution. After another washing step, all samples were mounted in FluorSave reagent (Merck Millipore #345789). GSC numbers were determined based on the morphology and position of their anteriorly anchored spherical spectrosome (Ables and Drummond-Barbosa, 2010; Ameku et al., 2018; Ameku and Niwa, 2016). Cap cells were identified by immunostaining with anti-LaminC antibody as previously described (Ables and Drummond-Barbosa, 2010).

### *Ex vivo* ovary culture

We used 5–6 day-old females. The ovaries were dissected in Schneider’s *Drosophila* medium (Thermo Fisher Scientific #21720024) and isolated from oviduct using forceps. Approximately 5–6 ovaries were immediately transferred to a dish containing 3 mL of Schneider’s *Drosophila* Medium supplemented with 15% fetal calf serum and 0.6% penicillin-streptomycin with/without the addition of OA (Sigma, final concentration of OA is 0–1000 mM) and 20E (Enzo Life Sciences; final concentration of 20 nM). The cultures were incubated at RT (except for *EcR* mutant flies, Figure 3D) for 16 h, and the samples were immunostained to determine the GSC number.

### *Ex vivo* calcium imaging

We employed the previously described imaging methods to visualize GSC behavior (Morris and Spradling, 2011; Reilein et al., 2018). For the live imaging, the ovaries dissected from adult virgin female flies were placed on a glass bottom dish (IWAKI #4970-041) with 3 mL of Schneider’s *Drosophila* Medium and 100 µL of the test reagent (Schneider’s *Drosophila* Medium containing 300 mM OA) placed directly at the center of each dish. The images were obtained with a × 40 objective lens (water-immersion) using a Zeiss LSM 700 confocal microscope and were recorded every 4 sec. The GCaMP6s fluorescence intensity in the escort cell was then calculated for each time point. The ratio of fluorescence (ΔF) at each time point was calculated by normalizing the fluorescence with the initial fluorescence (F0). The initial fluorescence (F0) is an average GCaMP6s fluorescence intensity before adding the test reagent.

### Optogenetic activation of the escort cells

Red-shifted channelrhodopsin CsChrimson (Klapoetke et al., 2014) was used to increase the [Ca^2+^]_i_ in the escort cells by light. *UAS-CsChrimson* was expressed using *c587-GAL4* with *nSyb-GAL80*. All crosses and the early development of flies were performed under dark conditions. The experiment was done at 25 °C. Adult flies were raised with standard food for 3 d after eclosion and then with standard food with 1 mM all-trans-retinal (ATR) for 3 d. Subsequently, they were kept in the presence of orange–red light from LED for 24 h. LED light was shined from the outside of the plastic chamber covered by aluminum foil to enhance light intensity.

### Statistical analysis

All experiments were performed independently at least twice. Fluorescence intensity in confocal sections was measured via ImageJ. For pMad quantification, signal intensity was calculated by measuring the fluorescence intensity in GSCs and CBs, which were co-stained with anti-Vasa antibody to visualize their cell boundaries. Sample sizes were chosen based on the number of independent experiments required for statistical significance and technical feasibility. The experiments were not randomized, and the investigators were not blinded. All statistical analyses were carried out using the “R” software environment. The P value is provided in comparison with the control and indicated as * for P ≤ 0.05, ** for P ≤ 0.01, *** for P ≤ 0.001, and “NS” for non-significant (P > 0.05).

## Acknowledgements

We thank Toshiro Aigaki, Aki Ejima, Kyung-An Han, Yukako Hattori, Yoshiki Hayashi, Makoto Ihara, Young-Joon Kim, Satoru Kobayashi, Kazuhiko Matsuda, Masayuki Miura, Akira Nakamura, Takashi Nishimura, Andrea Page-McCaw, Nobert Perrimon, Hiroko Sano, Jianjun Sun, Nobuaki Tanaka, James W. Truman, Tadashi Uemura, Daisuke Yamamoto, the Bloomington Stock Center, the Kyoto Stock Center (DGRC), the National Institute of Genetics, the Vienna Drosophila Resource Center, and the Developmental Studies Hybridoma Bank for providing stocks and reagents; Aki Hori and Reiko Kise for their technical assistance; and Satoru Kobayashi and Shosei Yoshida for their helpful discussion. Y.Y. and T.A. were recipients of the fellowship from the Japan Society for the Promotion of Science. This work was supported by grants from KAKENHI (26250001 and A17H01378 to H.T., 18J20572 to Y.Y., and 15J00652 to T.A), AMED-PRIME, AMED (17gm6010011h0001 to T.K.), AMED-CREST, AMED (19gm1110001h0003 to R.N.), and the Takeda Science Foundation to R.N.

---

**Figure 1-figure supplement 1.**
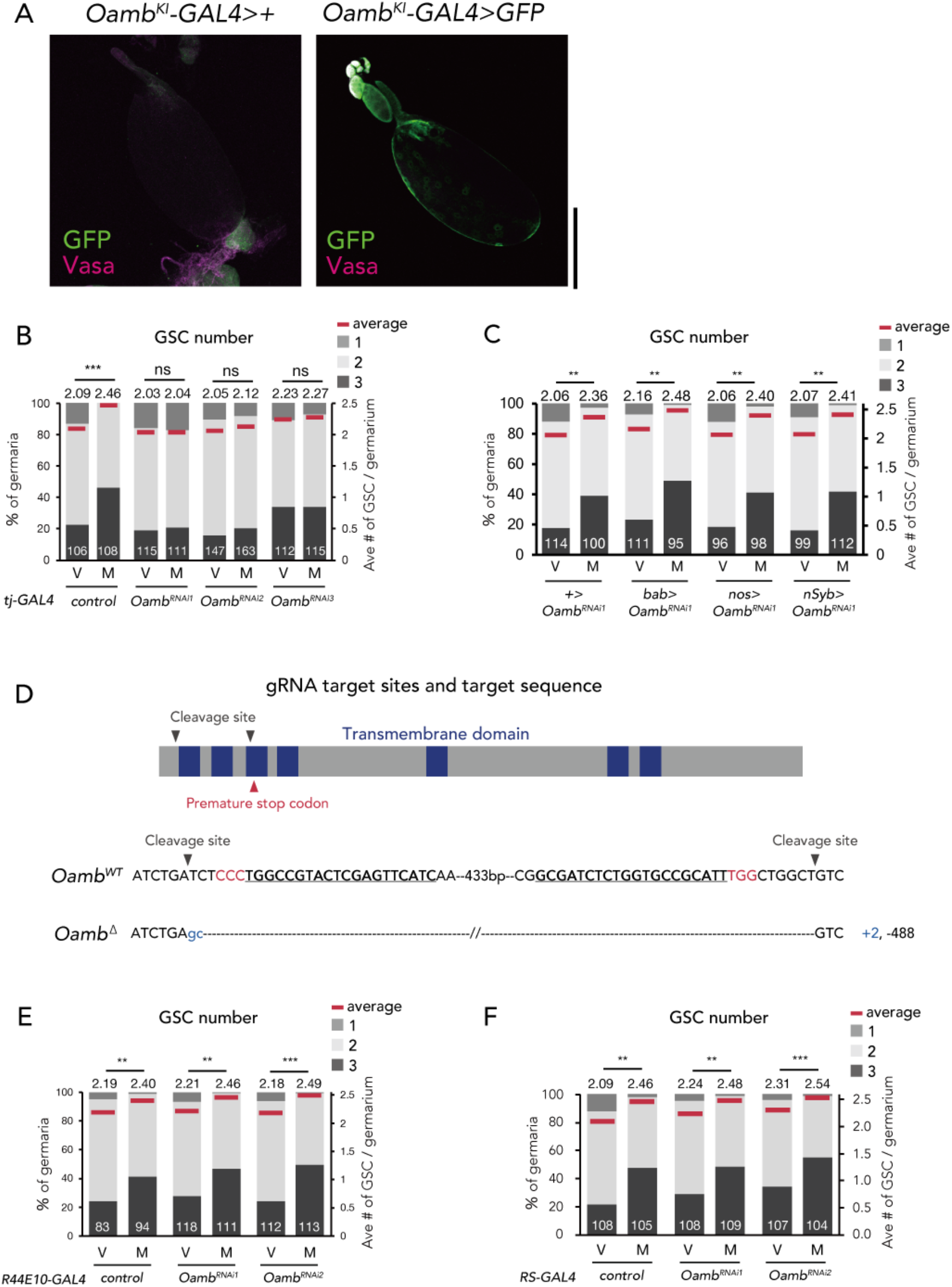
Oamb acts in the escort cells for post-mating GSC increase. (A) Immunofluorescence of stage-14 egg chamber expressing the *20xUAS-6xGFP* reporter under *Oamb^KI^-Gal4* (right). *GFP* expression was not observed in the control (*Oamb^KI^>+*; left). Scale bar, 200 µm. (B-C and E-F) Frequencies of germaria containing 1, 2, and 3 GSCs (left vertical axis) and the average number of GSCs per germarium (right vertical axis) in virgin (V) and mated (M) female flies in (B) *Oamb* RNAi by *tj-GAL4*; (C) *Oamb* RNAi in cap cells (by *bab-GAL4*), nervous system (by *nSyb-GAL4*), and germ cells (by *nos-GAL4*); (E) *Oamb* RNAi in mature follicle cells by (*R44E10-GAL4*); and (F) *Oamb* RNAi in the oviduct (by *RS-GAL4*). The number of germaria analyzed is indicated inside the bars. (D) A schematic representation of gRNA target sites (cleavage sites: grey arrowhead) and premature stop codon (red arrowhead) in coding sequences of *Oamb* genes. Regions of the putative transmembrane domains of Oamb are highlighted in violet. The target locus in Cas9-induced mutant was PCR-amplified and sequenced. The WT sequence is shown on the top of sequences as reference. The Cas9-gRNA target sequence is underlined with the PAM indicated in red. Inserted nucleotides are indicated in blue lowercase letters. The indel size is shown next to the sequence. The indel mutation results in a premature stop codon. Wilcoxon rank sum test with Holm’s correction was used for B, C, E, and F. ***P ≤ 0.001 and **P ≤ 0.01; NS, non-significant (P > 0.05).

**Figure 2-figure supplement 1.**
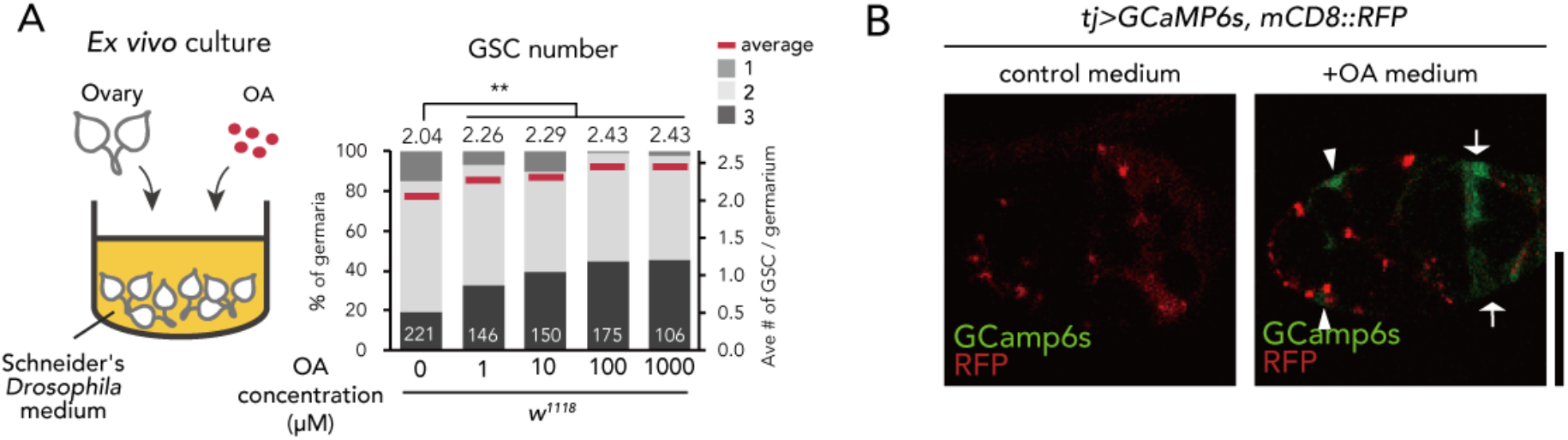
OA treatment induces GSC increase. (A) Frequencies of germaria containing 1, 2, and 3 GSCs (left vertical axis) and the average number of GSCs per germarium (right vertical axis) in virgin female flies. The addition of OA to the medium is sufficient to induce GSC increase. (B) Representative images of adult female germaria in response to OA in *tj>GCaMP6s; mCD8::RFP*. Note that calcium response was observed in the escort cells (arrowheads) and follicle cells (arrow) of the germarium. Scale bar, 10 µm. Wilcoxon rank sum test with Holm’s correction was used for statistical analysis. ***P ≤ 0.001.

**Figure 4-figure supplement 1.**
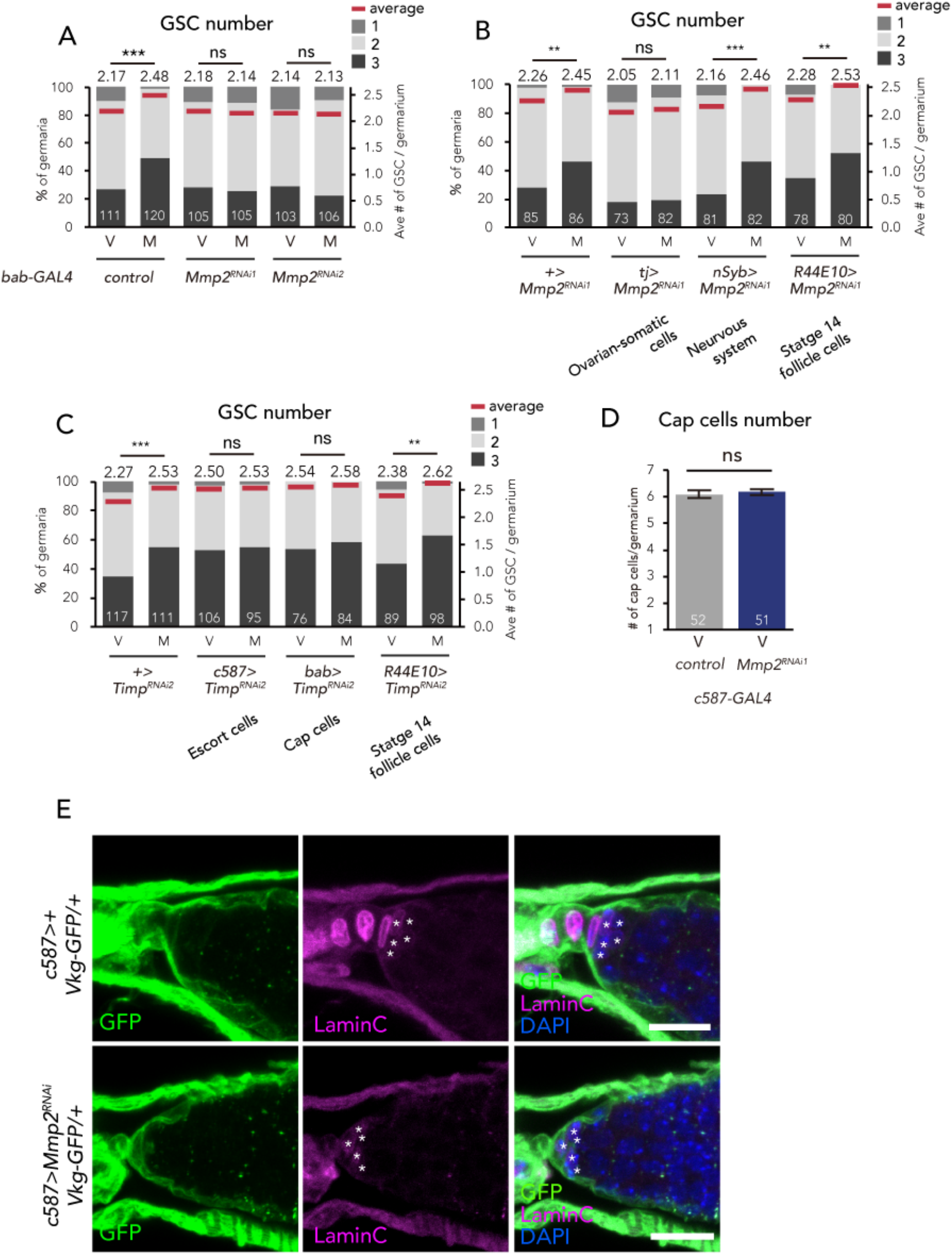
Mmp2 is necessary in the escort cells to induce GSC increase. (A-C) Frequencies of germaria containing 1, 2, and 3 GSCs (left vertical axis) and the average number of GSCs per germarium (right vertical axis) in virgin (V) female flies. *bab>+* flies were used as the control in A. *Mmp2* RNAi in the (A) cap cells by *bab-GAL4* driver and in the (B) nervous system by *nSyb-GAL4*, mature follicle cells by *R44E10-GAL4*, and ovarian somatic cells by *tj-GAL4*. (C) *Timp* RNAi in the escort cells by *c587-GAL4*, cap cells by *bab-GAL4*, and mature follicle cells by *R44E10*-GAL4. (D) The number of cap cells in the control and *Mmp2* RNAi driven by *c587-GAL4*. Values on y-axis are presented as the mean with standard error of the mean. (E) Representative images of *Vkg::GFP* adult female germaria immunostained with anti-GFP antibody (green), anti-LaminC antibody (red), and DAPI. The cap cells are indicated with asterisk. Note that the Vkg::GFP signal around cap cells is not affected even in *Mmp2* RNAi flies. Scale bar, 10 µm. Wilcoxon rank sum test with Holm’s correction was used for A, B, C and D. ***P ≤ 0.001 and **P ≤ 0.01; NS, non-significant (P > 0.05).

**Figure 5-figure supplement 1.**
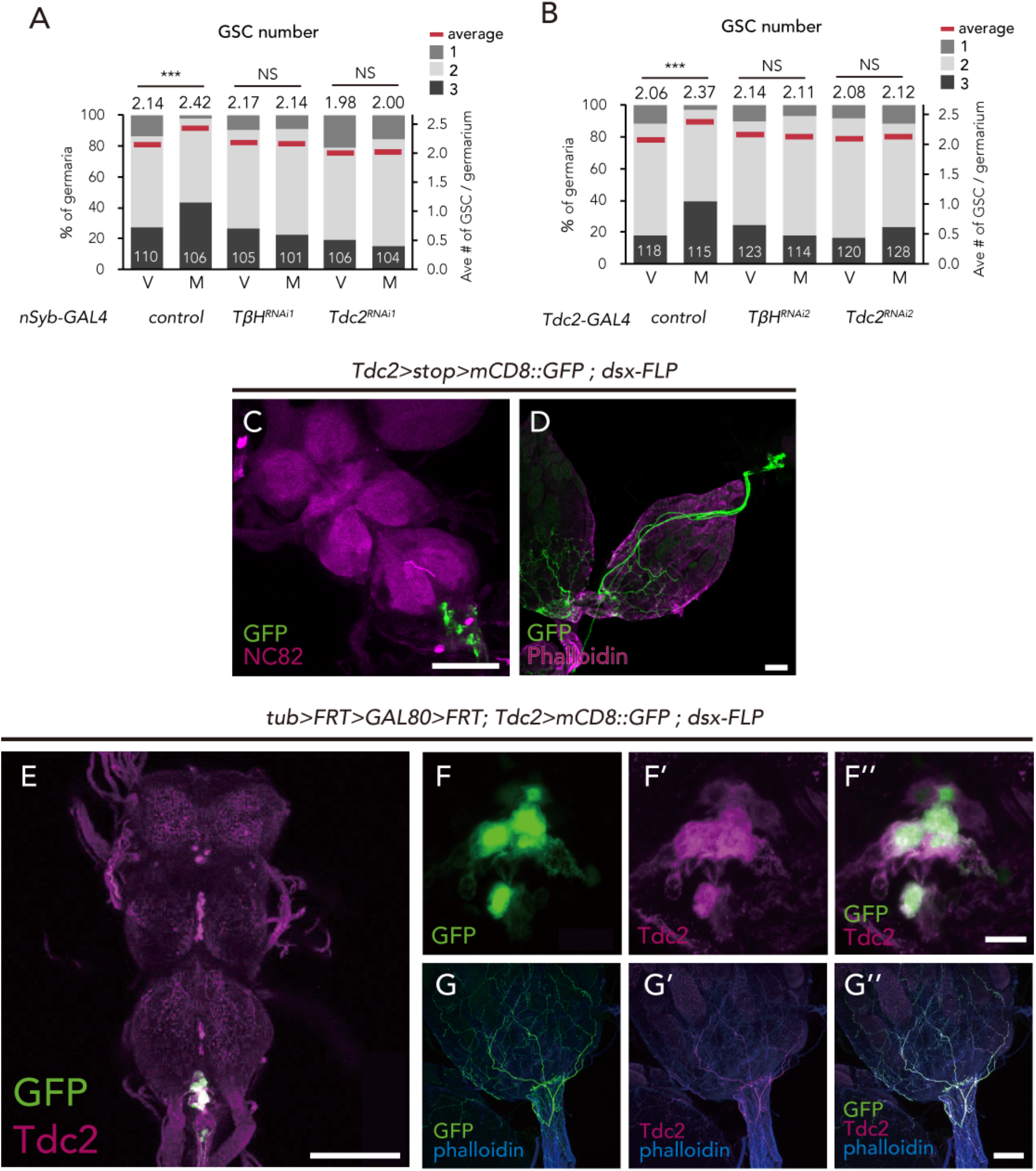
*dsx^+^ Tdc2^+^* neurons control GSC increase. (A-B) Frequencies of germaria containing 1, 2, and 3 GSCs (left vertical axis) and the average number of GSCs per germarium (right vertical axis) in virgin (V) and mated (M) female flies using *nSyb-GAL4* (A) and *Tdc2-GAL4* (B). (C-D) Images of *dsx^+^ Tdc2^+^* neurons expressing the *UAS>stop>mCD8::GFP* reporter under *Tdc2-Gal4* with *dsx-FLP*. *GFP* expression was detected only in the abdominal ganglion neurons projecting to the ovary. Images of the abdominal ganglion and the reproductive system are shown in C and D, respectively. (E-G) Immunofluorescence of *dsx^+^ Tdc2^+^* neurons expressing *UAS>mCD8::GFP* reporter under *Tdc2-Gal4* with *tub>FRT>GAL80>FRT>*. *GFP* expression was only observed in the anti-Tdc2 positive neurons in the abdominal ganglion that projected to the ovary (E). Scale bars, 100 µm in C, D, E, and G; 10 µm in F. Wilcoxon rank sum test with Holm’s correction was used. ***P ≤ 0.001; NS, non-significant (P > 0.05).

**Figure 6-figure supplement 1.**
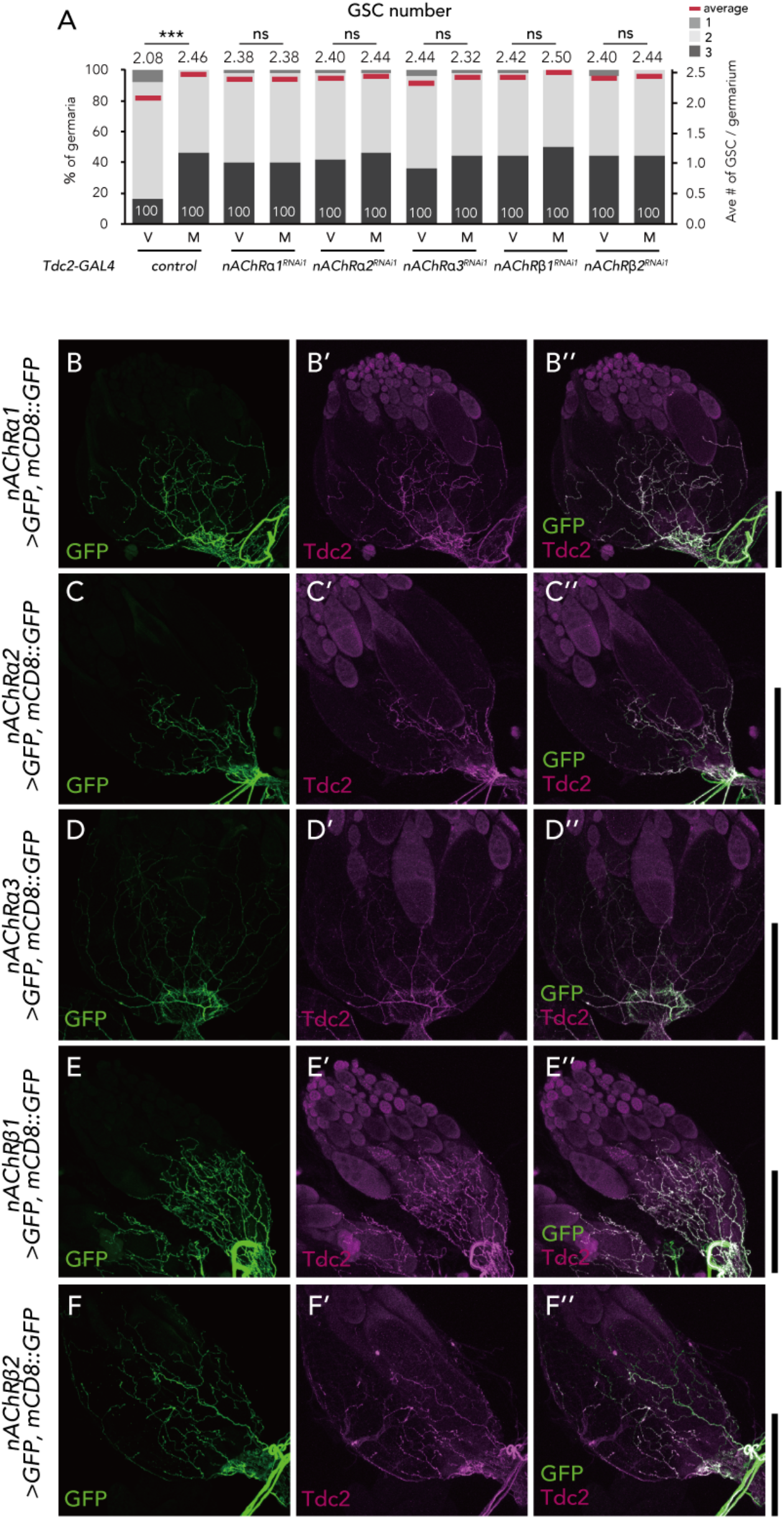
*nAChRs* are expressed in the ovary-projecting *Tdc2* neurons. (A) Frequencies of germaria containing 1, 2, and 3 GSCs (left vertical axis) and the average number of GSCs per germarium (right vertical axis) in virgin (v) female flies. (B-F) Representative images of the ovaries stained by anti-GFP (B-F) and anti-Tdc2 (B’-F’). Both signals merged on the surface of the ovary (B’’-F’’). Scale bar, 50 µm. Wilcoxon rank sum test was used. ***P ≤ 0.001; NS, non-significant (P > 0.05).

**Figure 6-figure supplement 2.**
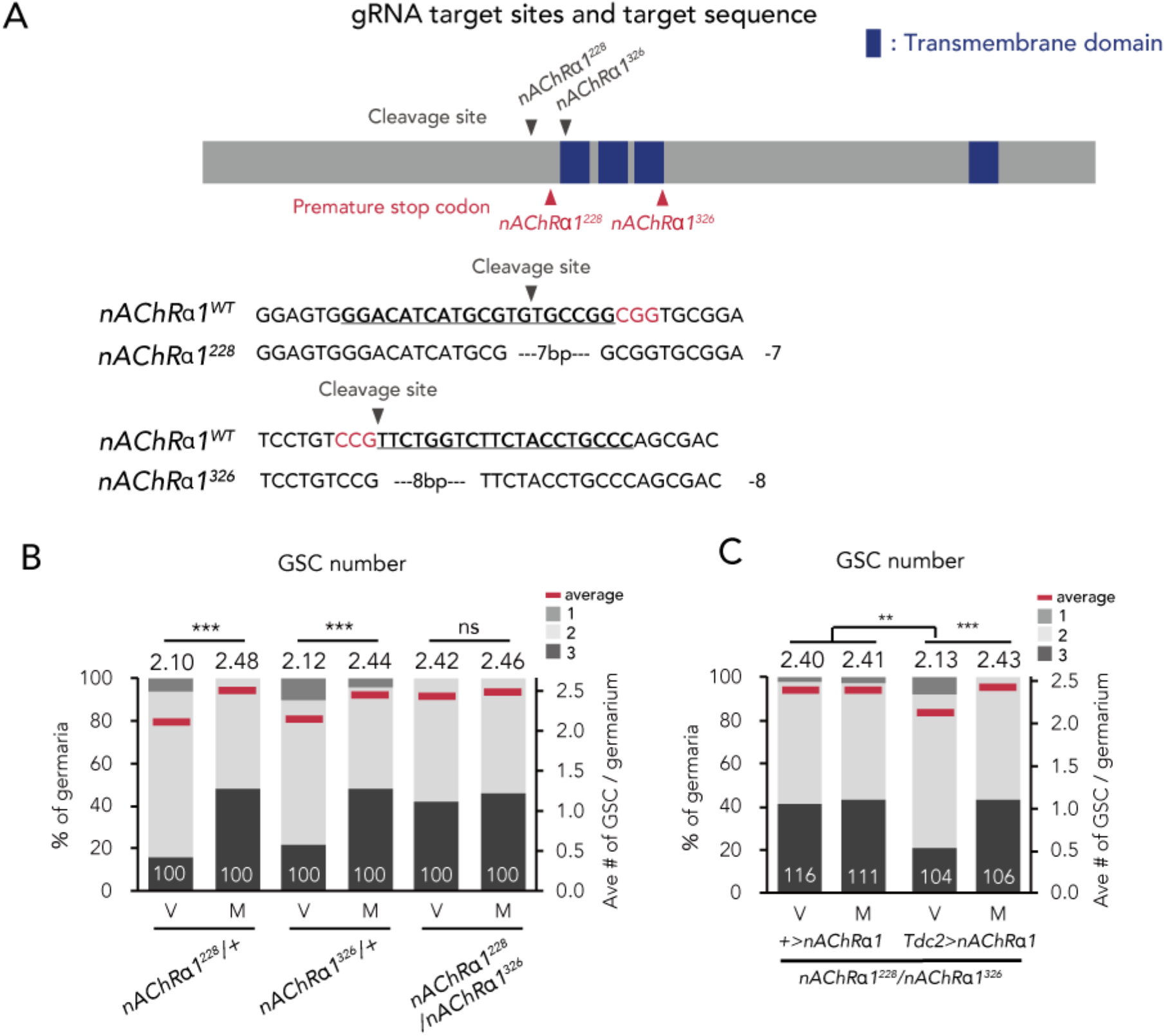
nAChRα1 in the *Tdc2* neurons regulates GSC increase. (A) Schematic representation of gRNA target sites (cleavage sites: grey arrowhead) and premature stop codon (red arrowhead) in coding sequences of *nAChRα1* genes. Regions of the putative transmembrane domains of *nAChRα1* are highlighted in violet. The target locus in Cas9-induced mutant was PCR-amplified and sequenced. The WT sequence (*nAChRα1^WT^*) is shown as reference. The Cas9-gRNA target sequence is underlined with the PAM indicated in red. The indel size is shown next to the sequence. The indel mutation results in a premature stop codon at the 228^th^ and 326^th^ amino acid sequence. (B-C) Frequencies of germaria containing 1, 2, and 3 GSCs (left vertical axis) and the average number of GSCs per germarium (right vertical axis) in virgin (V) and mated (M) female flies. (B) GSC numbers in *nAChRα1* genetic mutants. (C) *nAChRα1* overexpression in the *Tdc2* neurons was sufficient to restore increased GSC in *nAChRα1* mutants. Wilcoxon rank sum test with Holm’s correction was used. ***P ≤ 0.001 and **P ≤ 0.01; NS, non-significant (P > 0.05).

